# Discovery and characterization of chromosomal inversions in the arboviral vector mosquito *Aedes aegypti*

**DOI:** 10.1101/2024.02.16.580682

**Authors:** Jiangtao Liang, Noah Rose, Ilya I. Brusentsov, Varvara Lukyanchikova, Dmitry Karagodin, Yifan Feng, Andrey A. Yurchenko, Atashi Sharma, Massamba Sylla, Joel Lutomiah, Athanase Badolo, Ogechukwu Aribodor, Diego Ayala, Cassandra Gonzalez Acosta, Barry Wilmer Alto, Nazni Wasi Ahmad, Teoh Guat Ney, Zhijian Tu, Andrea Gloria-Soria, William C. Black, Jeffrey R. Powell, Igor V. Sharakhov, Carolyn S. McBride, Maria V. Sharakhova

## Abstract

Chromosomal inversions play a fundamental role in evolution and have been shown to regulate epidemiologically important traits in malaria mosquitoes. However, they have never been characterized in *Aedes aegypti*, the major vector of arboviruses, because of the poor structure of its polytene chromosomes. In this study, we applied a Hi-C proximity ligation approach to identify chromosomal inversions in 25 strains of *Ae. aegypti*, acquired from its worldwide distribution, as well as in one strain of *Ae. mascarensis*. The study identified 21 multi-megabase inversions with uneven distributions along the three chromosomes. All chromosomal inversions, including one specific for *Ae. mascarensis*, were polymorphic. Nevertheless, geographic origin separated the strains into two clusters carrying African and non-African inversions suggesting their potential association with *Ae. aegypti* subspecies. Some of the inversions colocalized with chemoreceptor genes and quantitative trait loci associated with pathogen infection, implicating the potential role of inversions in host choice and disease transmission.

---

Chromosomal inversions are important players in the evolution of diverse organisms including plants (Huang and Rieseberg 2020), vertebrate animals (Ferguson-smith and Trifonov 2007; Kosuthova and Solc 2023; Oliveira da Silva *et al*. 2024), and insects (Ayala and Coluzzi 2005; Fuller *et al*. 2019; Jay and Joron 2022). They represent balanced genome rearrangements in which a piece of the chromosome is flipped 180 degrees, producing a reverse order of the chromosomal region without gain or loss of the genetic material. In many organisms, at meiosis, heterozygote crossovers in the inverted region produce inviable gametes with acentric fragments and dicentric chromatids, thus, reducing or eliminating recombination between the breakpoints. As a result of reduced or eliminated recombination, an inversion is inherited intact as a “supergene” containing multiple divergent alleles (Thompson and Jiggins 2014; Berdan *et al*. 2022; Berdan *et al*. 2023). These supergenes often have dramatic phenotypic effects, much greater than single genes. Following pioneering studies by T. Dobzhansky in the 1930s on polytene chromosomes in *Drosophila* (Krimbas and Powell 1992), chromosomal inversions were used to study evolutionary genetics in other Dipteran species with well-developed polytene chromosomes, such as malaria mosquitoes (Kitzmiller 1976; Stegniy 1991; Coluzzi *et al*. 2002). Inversions have been shown to modulate epidemiologically important phenotypes in the major African malaria vector *Anopheles gambiae* (Lanzaro *et al*. 1998; Coluzzi *et al*. 2002; Gray *et al*. 2009; Simard *et al*. 2009; Ayala *et al*. 2014; Ayala *et al*. 2017; Riehle *et al*. 2017). However, identification of chromosomal inversions in aedini mosquitoes has been challenging because their large and highly repetitive genomes (Matthews *et al*. 2018) result in poorly developed polytene chromosomes that cannot be used for cytogenetic analyses (Campos *et al*. 2003). Previous genetic mapping experiments provided indirect evidence of chromosomal inversions in *Ae. aegypti* from Senegal (Bernhardt *et al*. 2009). Two putative chromosomal rearrangements have also been identified in Senegalese strains of *Ae. aegypti* using fluorescence *in situ* hybridization (FISH) (Dickson *et al*. 2016). More recently, linked-read genomic sequencing suggested the presence of multiple micro-inversions in 9 strains of *Ae. aegypti* (Redmond *et al*. 2020). Nevertheless, inversion polymorphism in natural populations of *Ae. aegypti* remains almost completely unknown and requires further study.

The *Ae. aegypti* mosquito transmits numerous arboviral infections with enormous impact on human health (Powell 2016b). This species was originally recognized as a major vector of yellow fever virus, which devastated human populations in the New World between the 17^th^ and early 20^th^ century when an effective vaccine was developed (Powell 2016a; Fijman and Yee 2022). Since the 1950s, another disease transmitted by *Ae. aegypti*, dengue fever, rapidly spread across the globe and became the leading arboviral disease of the 21^st^ century (Gubler 2012). Dengue is now endemic in 100 countries and has had an 8-fold increase in incidence over the last two decades (2022). Chikungunya and Zika fevers are two other diseases of rising concern that are transmitted by *Ae. aegypti* (Nabel and Zerhouni 2016; Powell 2016a). Zika virus caused an epidemic outbreak in Brazil in 2015 with 1.5 million cases and was shown to be related to more than 5 thousand cases of uncommon microcephaly in newborns (Faria *et al*. 2016; Weaver *et al*. 2016). Because *Ae. aegypti* is distributed across the global tropics and subtropics and thrives in human environments (Powell 2016a), the diseases associated with this mosquito currently threaten half of the world’s human population (Olagnier *et al*. 2016; 2022). This raises concerns about the risk of new epidemics in the future (Fontenille and Powell 2020). Better understanding of the genetic diversity in this species is critical for implementing effective vector control efforts.

Due to its wide geographical distribution, *Ae. aegypti* is genetically and phenotypically diverse, consisting of two subspecies *Ae. aegypti aegypti* (*Aaa*) and *Ae. aegypti formosus* (*Aaf*), which differ from each other in body coloration, behavior, habitats, association with humans, and ability to transmit pathogens (Mattingly 1957; Sylla *et al*. 2009; McBride *et al*. 2014; Powell *et al*. 2018; Aubry *et al*. 2020). The ancestral *Aaf* subspecies is a dark, sylvan form found in sub-Saharan Africa in various environments, including forest, rural, suburban, and urban settings (Powell and Tabachnick 2013; Tabachnick 2013; McBride *et al*. 2014; Rose *et al*. 2020). In contrast, the derived *Aaa* subspecies is a lighter form mostly found outside of Africa and is strongly associated with human environments. It has been proposed that the human associated lineage arose in Africa in response to the extremely long and hot dry seasons in the Sahel region of West Africa around 5000 years ago when the Sahara dried and water stored by humans became the major aquatic niche available for mosquitoes (Powell *et al*. 2018; Rose *et al*. 2020; Rose *et al*. 2023). This human-associated “proto-*Aaa*” lineage is believed to have then migrated to the New World on ships during the trans-Atlantic slave trade in the 1500s (Brown *et al*. 2011; Powell and Tabachnick 2013; Brown *et al*. 2014; Gloria-soria *et al*. 2016; Powell *et al*. 2018). While genetic, behavioral, and ecological data strongly suggest that introduction to the New World was from West Africa, it is less clear from exactly where in West Africa. Present day Angola was the site of departure of the highest number of slave ships and genetic data are consistent with this being the proximate origin of New World populations (Kotsakiozi *et al*. 2018). Interestingly, a second, more recent behavioral shift towards human-biting appears to be happening in response to rapid urbanization in West Africa over the last 20–40 years (Rose *et al*. 2020; Rose *et al*. 2023). Chromosomal inversions may contribute to the differentiation between the *Ae. aegypti* subspecies, particularly to adaptation of this mosquito to human hosts and habitats.

The *Ae. aegypti* subspecies display striking differences in their blood-feeding preferences (McBride 2016) and susceptibilities to different arboviral infections (Souza-net *et al*. 2019). While *Aaf* exhibits opportunistic blood choice and its feeding behavior likely depends on host availability, *Aaa* is strongly anthropophilic. Interestingly, genome comparisons of multiple strains of *Ae. aegypti* with different origins identified several key chromosomal regions associated with specialization to humans (Rose *et al*. 2020). Studies on vector competence found that *Aaa* is a better vector of arboviruses than *Aaf* (Lorenz *et al*. 1984; Tabachnick *et al*. 1985; Wallis *et al*. 1985; Sylla *et al*. 2009; Aubry *et al*. 2020), although there are many exceptions (Sylla *et al*. 2009; Dickson *et al*. 2014). Even though genetic mapping of quantitative trait loci (QTL) is not precise and provides only approximate locations of genetic factors linked to the particular phenotypes, an integration of physical and linkage maps indicated the presence of 5 chromosomal clusters of QTL related to different infections, suggesting that the transmission of different pathogens may be controlled by the same genomic loci (Timoshevskiy *et al*. 2013). Similar to *An. gambiae* (Lanzaro *et al*. 1998; Coluzzi *et al*. 2002; Gray *et al*. 2009; Simard *et al*. 2009; Ayala *et al*. 2014; Ayala *et al*. 2017; Riehle *et al*. 2017), chromosomal inversions can be related to the development of epidemiologically important phenotypes in *Ae. aegypti* mosquitoes.

The availability of a high-quality reference genome for *Ae. aegypti* (Matthews *et al*. 2018) now provides an opportunity to use genome-based technologies to detect chromosomal inversions in natural populations of this mosquito. One suitable method is Hi-C proximity ligation. Unlike microscopy, this approach belongs to the molecular-based 3C (Chromosome, Conformation, Capture)-group of methods (Dekker *et al*. 2002) and provides insights into the 3D-organization of whole genomes (Lieberman-aiden *et al*. 2009; van Berkum *et al*. 2010). The presence of specific inter- and intra-chromosomal interactions and chromosomal rearrangements, such as deletions, inversions, and translocations, can be estimated by measuring the frequency of physical contacts between specific genomic loci in the nuclear space (Engreitz *et al*. 2012; Chakraborty and Ay 2017; Harewood *et al*. 2017). The specially developed Hi-C software, Juicebox (Durand *et al*. 2016; Robinson *et al*. 2018), provides visualization of different types of interactions on Hi-C heat-map plots. Hi-C maps display short-distance interactions as a clear diagonal line, while long-distance interactions appear as bright spots off the diagonal. In cases where an inversion occurs, the areas that interact with each other in the nuclear space shift off the diagonal line. This method successfully identified previously known inversions in *An. coluzzii* and *An. stephensi* (Lukyanchikova *et al*. 2022b) and helped to unambiguously identify inversion breakpoints in *An. gambiae* (Corbett-detig *et al*. 2019). This method also revealed two previously unknown chromosomal inversions in *An. atroparvus* and *An. merus* (Lukyanchikova *et al*. 2022b).

The observed abundance of polymorphic inversions in *Anopheles* mosquitoes suggested an intriguing possibility that inversions on homologous chromosomal arms of different species capture similar sets of genes and, thus, confer similar ecological adaptations. Indeed, it has been demonstrated that polymorphic inversions on the 2R arm of *An. gambiae* and on the homologous arms of *An. funestus* and *An. stephensi*, which are involved in environmental adaptations in these three species, nonrandomly share orthologous genes (Sharakhova *et al*. 2011b). However, it is unknown how far such homologies could extend phylogenetically as polymorphic inversions have not been characterized in mosquitoes outside of the *Anopheles* genus.

In this study, we used the Hi-C proximity ligation method to search for chromosomal inversions in a panel of *Ae. aegypti* strains representing much of their worldwide distribution as well as widely used laboratory colonies and a closely related species, *Ae. mascarensis*. We found multiple large inversions, confirmed their existence by FISH, characterized their geographic distribution, analyzed the homology of inversions between *Ae. aegypti* and *An. gambiae*, and tested for overlap with regions of the genome previously shown to be associated with olfaction and vector competence. Our work provides insight into the potential role of chromosomal rearrangements in *Ae. aegypti* behavior and vectorial capacity and builds a foundation for future research in these fields.

## Materials and Methods

### Mosquito colonies

Information about mosquito colonies and collections is summarized in Table S1 (Severson *et al*. 2002; Garcia-luna *et al*. 2018; Matthews *et al*. 2018; Rose *et al*. 2020; Metz *et al*. 2023). The study included 23 strains of *Ae. aegypti*, which were recently colonized. Of these, 12 strains were from 6 African countries across the species’ native range (Senegal, Burkina Faso, Nigeria, Gabon, Uganda, and Kenya), while 11 strains were from 5 non-African countries (Mexico, USA, Brazil, Thailand, and Malaysia). In addition, two old laboratory strains of *Ae. aegypti* (Liverpool and inbred Liverpool strain, RU3) and *Ae. mascarensis*, a closely related species of *Ae. aegypti*, from Mauritius Island, were also utilized. Colonies were kept under standard conditions in incubators (Benedict and Dotson 2015). Mosquito eggs were hatched in distilled water and kept at 27 °C until the larval and pupal stages in an incubator. After emerging from pupae, the adults were maintained at the same temperature with a 70 ± 5% relative humidity and a 12-hour cycle of light and darkness. Larvae and adults were fed with fish food and 10% sugar water, respectively. 5–7-day-old adult females were blood-fed using artificial blood feeders with defibrinated sheep’s blood (HemoStat Laboratories Inc., CA, USA). Egg dishes were placed in cages for oviposition approximately two to three days after blood feeding.

### Hi-C inversion detection

To discover chromosomal inversions, 2,235 adults from 20 strains and >6,000 embryos from 6 strains were used to build 49 Hi-C libraries (Table S1). Two Hi-C libraries were made for the majority of strains, except for three strains (MONT, GUER, and RU3), for which only one library each was constructed. We first utilized the *in situ* Hi-C protocol for library preparation that was developed for mosquito embryos and described previously (Lukyanchikova *et al*. 2022a). Each replicate was made from more than 1,000 embryos. We applied the Arima-HiC+ High Coverage kit (Arima Genomics, Carlsbad, CA, USA) for the adult stage of mosquitoes. The majority of the adults used here were males, which was needed for the equal capturing of both sex-determining chromosome 1, with exception for two Mexican strains (GUER and MONT) where both sexes were included due to the low amount of field collected materials. Adult mosquitoes were 1–3-day old. Hi-C libraries were prepared according to the protocols provided by the company (Arima Genomics, Carlsbad, CA, USA). The libraries were sequenced on the Illumina platform to get an approximate 50× coverage for each strain. For Hi-C heatmap development, sequence quality assessment was performed using FastQC software v.0.12.1 (Andrews 2010). The reads were aligned to the AaegL5 genome (Matthews *et al*. 2018) and Juicer Pipeline (Durand *et al*. 2016) was used to build the contact matrices and Hi-C heatmaps. A Juicebox software (Durand *et al*. 2016) was employed to visualize and compare heatmaps. Inversion identification was performed based on visual inspection of Hi-C maps as was done previously (Lukyanchikova *et al*. 2022b). Chromatin contact maps were built for each *Ae. aegypti* strain and inspected for long-range interactions. Long-distance contacts with a butterfly-like pattern were considered indicative of inversions.

### Probe preparation and fluorescent *in situ* hybridization (FISH)

The probes for FISH were designed based on gene exons or cDNA fragments of ≈3.5 kb sizes inside of the inversions according to breakpoints predicted by the Hi-C maps. These fragments were synthesized by PCR using Phusion DNA polymerase (Thermo Fisher Scientific, MA, USA) according to the manufacturer’s recommendations. Genomic DNA or cDNA were used as the templates (Masri *et al*. 2021). Detailed probe information is shown in Table S2. For cDNA synthesis, total RNA was extracted from 2–3-day-old female adult mosquitoes using TRIzol Reagent (Thermo Fisher Scientific, MA, USA). The reverse transcription reaction was performed in a 20 μl reaction mixture with 5 μg total RNA, 1× reaction buffer, 0.5 mM dNTPs, 100 pmol oligo(dT)_15–18_ primer, and 40 U Maxima H Minus Reverse Transcriptase (Thermo Fisher Scientific, MA, USA), for 30 min at 50 °C. The resulting cDNA was stored at −20 °C and 1⁄50–1⁄100 portion per reaction was used for PCR. For cloning, PCR products were excised from the gel, purified using the QIAquick Gel Extraction Kit (QIAGEN, MA, USA), and treated with T4 Polynucleotide Kinase (Sibenzyme, Novosibirsk, Russia). pBluescript SK (+) plasmid was digested by the restriction endonuclease EcoR V (Sibenzyme, Novosibirsk, Russia), treated with Thermolabile Alkaline Phosphatase (Sibenzyme, Novosibirsk, Russia), and purified using the QIAquick Gel Extraction Kit (QIAGEN, MA, USA). Phosphorylated PCR products were ligated with prepared plasmids using T4 DNA ligase (Sibenzyme, Novosibirsk, Russia) and transformed to chemical competent *Escherichia coli* cell XL1-Blue. Successful clones were validated by Sanger sequencing.

Chromosomal preparations were followed as described elsewhere (Sharakhova *et al*. 2011a; Liang *et al*. 2022). Imaginal discs were dissected from 4^th^ instar larvae. Good chromosome preparations were selected for FISH under an Olympus CX41 microscope (Olympus America, Inc., Melville, NY, USA) and FISH was performed using standard protocols (Timoshevskiy *et al*. 2012; Sharakhova *et al*. 2019). Purified plasmids were labeled with Cy3- or Cy5-dUTP (Enzo Life Sciences Inc., Farmingdale, NY, USA) using a nick translation method. After FISH, slides were analyzed and documented using a Zeiss AXIO fluorescent microscope with an Axiocam 506 mono digital camera (Carl Zeiss AG, Oberkochen, Germany) and a Zeiss LSM 880 Confocal Microscope (Carl Zeiss Microimaging, Inc., Thornwood, NY, USA). Previously described methods were utilized for probe mapping (Sharakhova *et al*. 2019). The FISH data are summarized in Table S3.

### Identification of SNPs specific for 1pA and 3qG inversions

To identify putative inversion-diagnostic SNPs for the two most widespread inversions, 1pA and 3qG, principal components analysis (PCA) of 393 previously-published *Ae. aegypti* genomes from Africa (Rose *et al*. 2020) was utilized. A two-step process was applied to minimize the effects of population structure. First, a PCA was conducted on the target genomic regions in a sample of individuals from Southern Senegal (BTT, KED, MIN, and PKT populations), where Hi-C analyses indicated the inversions should be polymorphic. A scatter plot of the first two principal components was visually inspected to confirm a bimodal or trimodal distribution of points along the first principal component, corresponding to different inversion genotypes. A trimodal distribution including standard (wildtype) homozygotes, inversion heterozygotes, and inverted homozygotes is expected for common inversions. A bimodal distribution representing standard homozygotes and inversion heterozygotes is only expected for rare inversions not present in the homozygous state. Second, we examined loadings on the first principal component, and selected the set of SNPs with loadings close to the highest observed absolute value for further analysis across the full set of 393 individuals (79 SNPs for 1pA, 67 for 3qG), and the inversion genotype was estimated by taking the consensus genotype across these putatively inversion-linked SNPs. Putative diagnostic SNPs were selected based on 1) perfect or near-perfect (Pearson correlation >0.99) correlation with the putative karyotype and 2) distributions among populations corresponding to those identified from prior Hi-C analyses. Finally, 5 and 3 SNPs were chosen as diagnostics for inversions 1pA and 3qG, respectively (Table 1).

**Table 1.** Diagnostic SNPs for inversions 1pA and 3qG. SNP showing perfect correlation with putative karyotype shown in bold.

| Inversion | Inversion genome coordinates (Mb) | Genome coordinates for diagnostic SNPs (bp) |
| --- | --- | --- |
| 1pA | 1.2–16.5 | 16421358, 16443906, 16452822, 16452923, 16453689 |
| 3qG | 404.1–409.3 | **404258781**, 404651310, 408994400 |

### Cluster analysis of the 1pA and 3qG inversion frequencies

Hierarchical clustering analysis was performed using the frequencies of autosomal chromosomal inversions at the population level to infer and visualize the relationship between the species and populations using Package for R “stats” (R_Core_Team 2021). A total of 387 previously published whole-genome sequences (WGS) of individual mosquitoes from 31 locations from 7 countries in Africa and 2 countries outside of Africa (Rose *et al*. 2020) were analyzed to estimate frequencies of the two most common inversions, 1pA and 1qG. The final analysis included 26 locations because, in four different cases, nearby (<50 km) populations with low numbers of mosquitoes were merged (Table S4). The reliability of data clustering was assessed using the “pvclust” package for R (Suzuki and Shimodaira 2006). The data were resampled 100 times and the similarity of different clustering variants was assessed using methods published elsewhere (Shimodaira 2002; Shimodaira 2004). The average values of these similarities were used as an indicator of cluster stability.

### Colocalization of inversions with chemoreceptor genes and quantitative trait loci related to vector competence

Colocalization of inversions with chemoreceptor genes (Matthews *et al*. 2018) and QTL, associated with mosquito competence for key pathogens (Bosio *et al*. 2000; Severson *et al*. 2002; Timoshevskiy *et al*. 2013) was determined. First, the numbers of genes inside of the inversion were identified (denoted as *k*). Then, the statistical significance of increased density of odorant receptor (OR), gustatory receptor (GR), and ionotropic receptor (IR) genes and QTL inside of the inversions was tested against the uniform distribution of them along the chromosomes. P-values were calculated as the upper tails of binomial distributions, i.e. *P*(*X* ≥ *k*) = 1−*CDF*(*k*−1, *B*(n, p)) = 1− *I*_1−*p*_(*n*−*k*+1, *k*), where *p*=*l*/*L, CDF*(*k*−1, *B*(*n, p*)) is the cumulative distribution function of binomial distribution *B* with parameters *p* and *n, I* is the incomplete beta function, *l* is the inversion length, *L* is the chromosome length, and *n* is the number of genes on the chromosome, {*n, k, l, L*} ∈ ℤ. The correction for multiple P-values was calculated using the Benjamini-Hochberg method in the “stats” R package (Benjamini and Hochberg 1995) (Table S5). We applied the same approach to determine overlap between inversions and QTL associated with vector competence to different infections (Table S5).

In addition, the chemoreceptor genes were combined into evolutionary clusters to exclude the negative effect of the cluster organization of chemoreceptor genes in the mosquito genomes on the significance of the results. For this purpose, phylogenetic trees were constructed for chemoreceptor genes of *Ae. aegypti*, as described previously (Ryazansky 2023). Branches with closely related genes on these trees were identified and combined into clusters of evolutionary related and closely located genes (Table S6). The distribution of the genes was then tested for compliance with an uniform distribution using the Distribution-FitTest function from the Mathematica v. 13.3 program (Wolfram 2023). P-values were calculated as described previously considering each cluster of chemoreceptor genes as one gene (Table S7).

Finally, a permutation test was used to calculate non-random distribution of inversions in respect to chemoreceptor genes (Cabrera *et al*. 2012). A total of 100 circular chromosome permutations, in which a single random offset was drawn from a uniform distribution across the length of each chromosome, was added to all chemoreceptor positions within a given permutation. Then the modulo of the resulting position with the length of the chromosome was used to assign the new position of each chemoreceptor gene by “rotating” the position of all chemoreceptor genes together, preserving the relative distribution of chemoreceptor genes within the chromosome in each permutation. The number of chemoreceptor gene-inversion intersections in each permutation using *bedtools intersect* was then calculated.

### Overlapping inversions in the *Ae. Aegypti* and *An. gambiae* genomes

To determine if chromosomal inversions in *Ae. aegypti* and in *An. gambiae* captured similar sets of genes, we first identified single copy orthologs between the species using OrthoFinder software (Emms an Kelly 2019) with default settings. The genomic positions of the inversions were identified based on previously published data (Sharakhov *et al*. 2006; Coulibaly *et al*. 2007; George *et al*. 2010; Lobo *et al*. 2010) summarized in Table S8. The identified single-copy orthologs were overlapped with the genomic coordinates between two species using the BEDTools function *intersect-loj* (Quinlan 2014). To calculate the significance of pairwise inversion overlaps between species, we used a hypergeometric test based on the single-copy orthologs inside and outside of each inversion. The raw P-values were adjusted with the Benjamini-Hochberg method. Calculations were performed using the R package “GeneOverlap” (shenlab-sinai 2023). Circos software was used to visualize the common regions with inversion polymorphism among the species (Krzywinski *et al*. 2009).

## Results

### Inversion discovery using Hi-C approach

In this study, we used the Hi-C technology to discover chromosomal rearrangements in *Ae. aegypti*. To develop Hi-C libraries, we tested adults from 20 strains and embryos from 6 strains (Table S1, see Method section for details). The inversions were recognized based on a butterfly-like pattern outside of the major diagonal in the Hi-C heatmap (Fig. 1, Fig. S1). Fig. 1 shows examples of the Hi-C results for chromosomal inversions in chromosome 1 (1pA, 1qD, 1qE, 1qF, 1qG) and misassembly 2. The coordinates of the inversion breakpoints were determined based on the center position of the butterfly-like pattern at a resolution of 5 kb. The lengths of inversions were identified based on breakpoint coordinates (Table 2). For some widespread inversions, we observed variations in the shape and intensity of the butterfly-like pattern across strains, possibly related to the frequency of the inverted karyotype in the pool of sequenced individuals. For example, Fig. 1B illustrates how the pattern observed at the 1pA inversion varied from almost a complete disruption of the major diagonal and well-pronounced butterfly pattern in the UGA strain (top line) to the complete absence of inversion signal in the SHM strain (bottom line). Overall, a higher abundance of individuals with the inversion resulted in a stronger and more intense butterfly-like pattern outside of the diagonal (Fig. 1B). After evaluating the Hi-C maps of all the strains collected from the field, the old laboratory strains, and *Ae. mascarensis*, we concluded that, according to our criteria, none of the inversions resulted in a complete disruption of the major diagonal for all the examined strains. Therefore, we classify all the inversions discovered in this study as polymorphic.

**Table 2.**
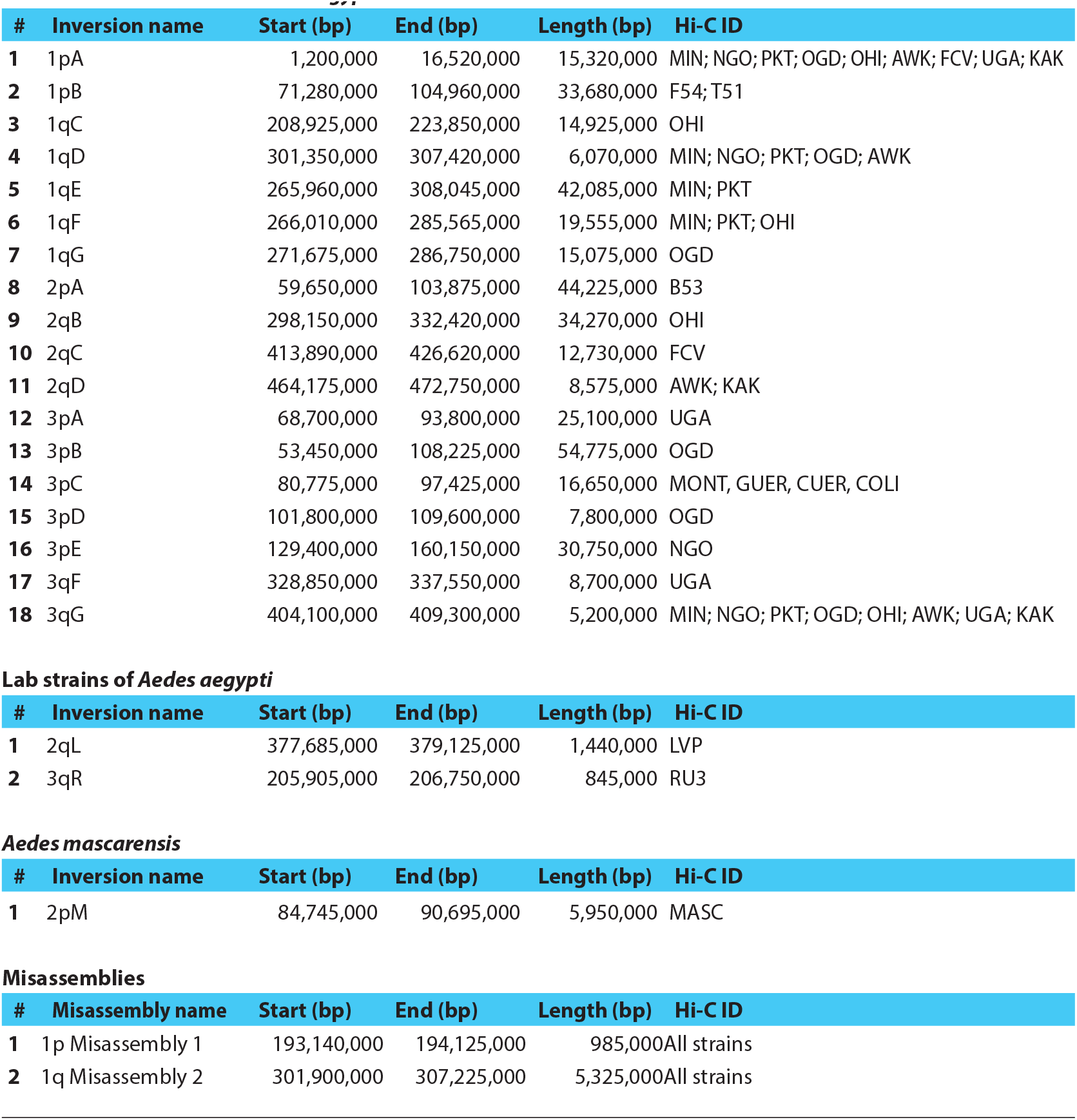
Chromosomal inversions in *Aedes aegypti* and *Aedes mascarensis* Field collected strains of *Aedes aegypti*.

| # | Inversion name | Start (bp) | End (bp) | Length (bp) | Hi-C ID |
| --- | --- | --- | --- | --- | --- |
| 1 | 1pA | 1,200,000 | 16,520,000 | 15,320,000 | MIN; NGO; PKT; OGD; OHI; AWK; FCV; UGA; KAK |
| 2 | 1pB | 71,280,000 | 104,960,000 | 33,680,000 | F54; T51 |
| 3 | 1qC | 208,925,000 | 223,850,000 | 14,925,000 | OHI |
| 4 | 1qD | 301,350,000 | 307,420,000 | 6,070,000 | MIN; NGO; PKT; OGD; AWK |
| 5 | 1qE | 265,960,000 | 308,045,000 | 42,085,000 | MIN; PKT |
| 6 | 1qF | 266,010,000 | 285,565,000 | 19,555,000 | MIN; PKT; OHI |
| 7 | 1qG | 271,675,000 | 286,750,000 | 15,075,000 | OGD |
| 8 | 2pA | 59,650,000 | 103,875,000 | 44,225,000 | B53 |
| 9 | 2qB | 298,150,000 | 332,420,000 | 34,270,000 | OHI |
| 10 | 2qC | 413,890,000 | 426,620,000 | 12,730,000 | FCV |
| 11 | 2qD | 464,175,000 | 472,750,000 | 8,575,000 | AWK; KAK |
| 12 | 3pA | 68,700,000 | 93,800,000 | 25,100,000 | UGA |
| 13 | 3pB | 53,450,000 | 108,225,000 | 54,775,000 | OGD |
| 14 | 3pC | 80,775,000 | 97,425,000 | 16,650,000 | MONT, GUER, CUER, COLI |
| 15 | 3pD | 101,800,000 | 109,600,000 | 7,800,000 | OGD |
| 16 | 3pE | 129,400,000 | 160,150,000 | 30,750,000 | NGO |
| 17 | 3qF | 328,850,000 | 337,550,000 | 8,700,000 | UGA |
| 18 | 3qG | 404,100,000 | 409,300,000 | 5,200,000 | MIN; NGO; PKT; OGD; OHI; AWK; UGA; KAK |

**Lab strains of *Aedes aegypti***
| # | Inversion name | Start (bp) | End (bp) | Length (bp) | Hi-C ID |
| --- | --- | --- | --- | --- | --- |
| 1 | 2qL | 377,685,000 | 379,125,000 | 1,440,000 | LVP |
| 2 | 3qR | 205,905,000 | 206,750,000 | 845,000 | RU3 |

***Aedes mascarensis***
| # | Inversion name | Start (bp) | End (bp) | Length (bp) | Hi-C ID |
| --- | --- | --- | --- | --- | --- |
| 1 | 2pM | 84,745,000 | 90,695,000 | 5,950,000 | MASC |

**Misassemblies**
| # | Misasassembly name | Start (bp) | End (bp) | Length (bp) | Hi-C ID |
| --- | --- | --- | --- | --- | --- |
| 1 | 1p Misasassembly 1 | 193,140,000 | 194,125,000 | 985,000 | All strains |
| 2 | 1q Misasassembly 2 | 301,900,000 | 307,225,000 | 5,325,000 | All strains |

**Fig. 1.**
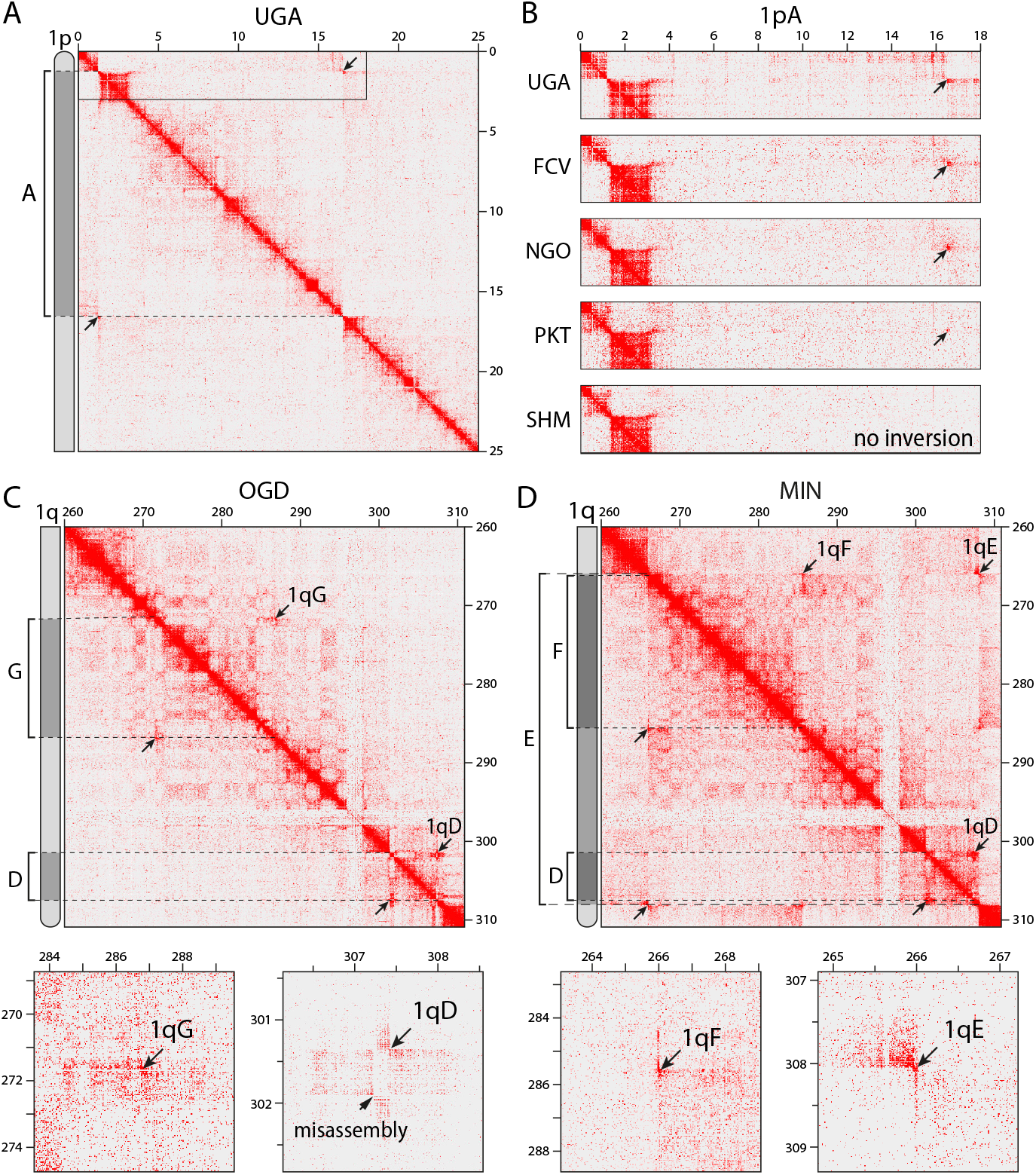
Inversion patterns in the Hi-C heat maps of the chromosome 1. **A)** Hi-C heat map for the 1pA inversion in the UGA strain. **B)** Different patterns of Hi-C heat maps associated with frequencies of the 1pA inversion in different strains. **C)** Hi-C heat map for the 1qG and 1qD inversions and misassembly 1 in the OGD strain. **D)** Hi-C heat map for three overlapping inversions, 1qD, 1qE, and 1qF inversions, in the MIN strain. A schematic representation of inversions indicated by brackets within the chromosome arms are shown on the left of panels A, C, and D. Arrows point to the center of the butterfly-like patterns. Break points are shown by dashed lines. Below the pictures on panels C and D are enlarged images of the inversions and misassembly 1. Chromosomal inversions are seen as a butterfly-like pattern outside of the major diagonal on the Hi-C heat maps. These patterns are more pronounced in strains with a higher abundance of individuals carrying inversion 1pA (panel B). Overlapping inversions 1qE and 1qF produce a disrupted butterfly pattern on the Hi-C heat map in panel D.

Strikingly, a puzzle of four overlapping inversions—1qD, 1qG, 1qE, and 1qF—was found in the telomere region of the 1q chromosome arm (Fig. 1C, D). Although inversions 1qD, 1qF, and 1qG were found as separate inversions in different strains (Table 2, Fig. 1C), inversion 1qE was always present together with 1qD and 1qF (e.g., in MIN and PKT, Senegal), representing an example of a triple inversion (Fig. 1D). This result suggests that inversion 1qE (42 Mb long) was originally formed on the chromosome that contained both of the smaller inversions 1qD (5 Mb long) and 1qF (19 Mb long) (Fig. 1C, D). As a consequence, inversions 1qE and 1qF formed a “broken butterfly-like” pattern in the Hi-C map of the MIN strain (Fig. 1D) because the proximal breakpoints of the 1qE and 1qF inversions were very close to each other (≈50 kb). Inversion 1qD in the same Hi-C map in the MIN strain formed a clear independent butterfly-shaped pattern because the distal breakpoints of inversions 1qD and 1qF were further from each other (≈625 kb). None of the described inversions shared breakpoints.

Finally, our study identified two misassemblies in the reference genome AaegL5.0 (Matthews *et al*. 2018). The lengths of misassemblies 1 and 2 were 0.98 Mb and 5.3 Mb, respectively. The presence of misassemblies were considered if the butterfly-like patterns outside of the major diagonal in Hi-C maps were determined in all examined strains (Table 2), including the RU3 strain, which was used for the genome assembly (Matthews *et al*. 2018). The larger misassembly 2 was located near inversion 1qD but had slightly different coordinates and was clearly differentiable in both the OGD and MIN strains in Fig. 1C and D, respectively.

Overall, based on Hi-C data, our study discovered 18 chromosomal inversions in recently colonized strains from around the world, 2 inversions in old laboratory strains, Liverpool (Severson *et al*. 2002) and the inbred derivative strain RU3 (Matthews *et al*. 2018), 1 inversion in *Ae. mascarensis*, and 2 potential misassemblies in the reference genome AaegL5.0 (Matthews *et al*. 2018).

### Inversion validation using fluorescence *in situ* hybridization

To validate the Hi-C results, chromosomal inversions were examined using FISH in mitotic chromosomes from imaginal discs of 4^th^ instar larvae. Six inversions (1pA, 1qD, 1qE, 1qF, 3pA, 3pB) and misassembly 2 were selected for validation based on their abundance, geographical distribution, and chromosomal location. Since some of these inversions, as well as misassembly 2, overlapped, we were able to perform FISH using only three pairs of probes: AAEL012233 and AAEL022285 probes for the 1pA inversion, AAEL025976 and AAEL023895 probes for the 1qD and 1qE inversions and misassembly 2 on the 1q arm, and AAEL007127 and AAEL019648 probes for the 3pA and 3pB inversions (Table S2).

A scheme and examples of FISH experiments in chromosome 1 are shown in Fig. 2. As each of the paired probes was labeled by a different fluorescent dye (Cy3 or Cy5), standard, inverted, and heterozygote karyotypes were identified by the order of the probe signals along the chromosomes based on their proximity to the telomere. For the 1pA inversion, we performed FISH for the RU3, PKT, and UGA strains. FISH results demonstrated that only individuals with standard arrangements were present in the RU3 and PKT strains but only inverted arrangements were found in all individuals of the UGA strain (Fig. 2, Table S3).

**Fig. 2.**
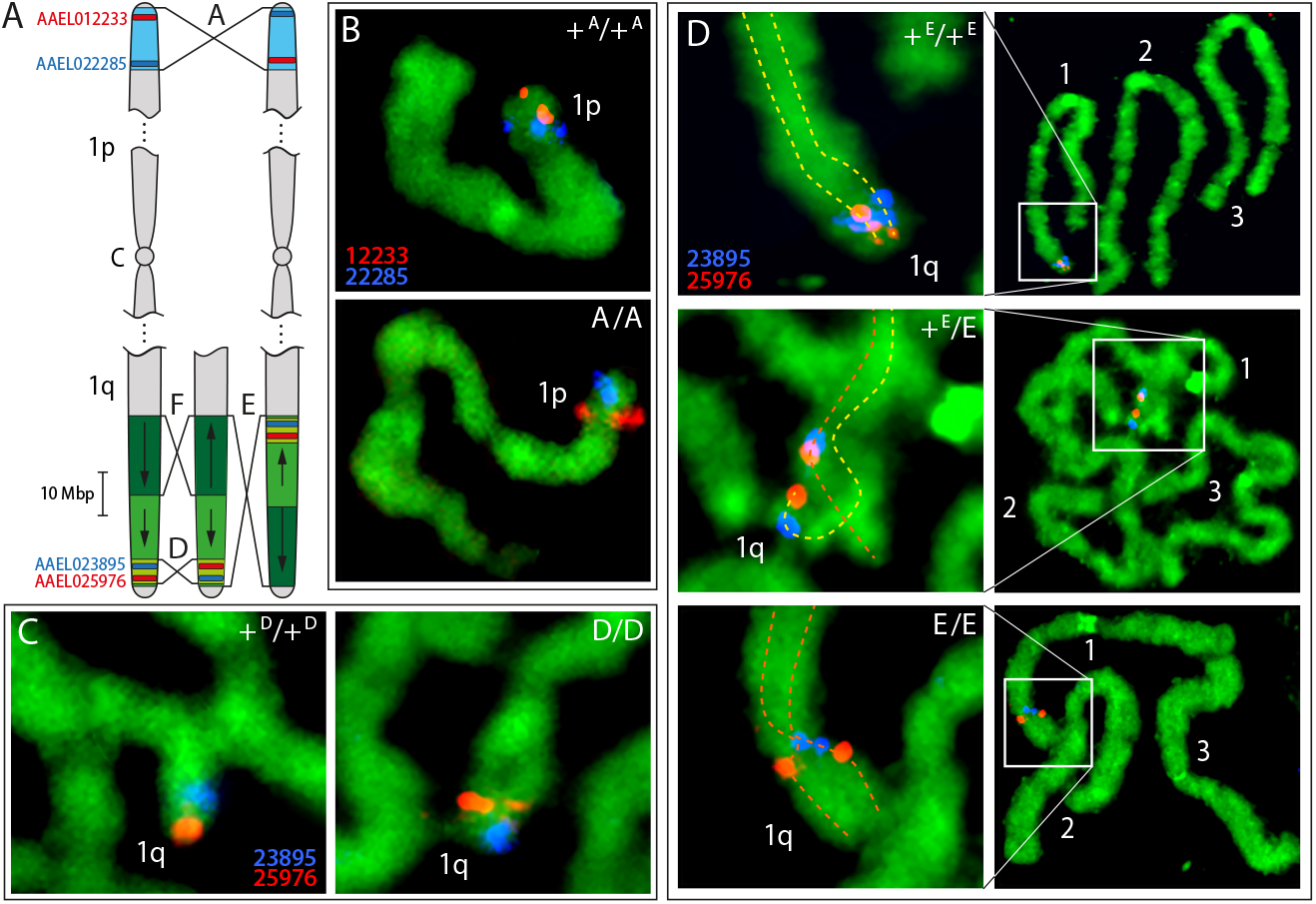
Validation of inversions in chromosome 1 using FISH. **A**. A diagram of FISH experiments for chromosome 1. Chromosomes are schematically shown by gray color. Letter C stands for the centromere. Inversion 1pA is shown in blue color, inversions 1qE, 1qD, and 1qF are shown by different shades of green color. Chromosome arms 1p and 1q on the left of the panel represent standard arrangements; chromosome arms in the middle and on the right are shown in inverted arrangements. Arrows inside the inversion regions of 1qE and 1qF indicate orientations of standard arrangements. Positions of probes used for FISH, AAEL012233, AAEL025976 and AAEL022285, AAEL023895, are indicated by red and blue colors, respectively. The corrected pattern of misassembly 2 is shown in Fig. S2. Breakpoint positions are indicated by brackets. **B–D**. Examples of FISH for validation of the inversions 1pA (B), 1qD (C), and 1qE (D). 1p and 1q stand for the chromosome arms. Numbers 1, 2, and 3 stand for the chromosome numbers. All images represent chromosomes at the prometaphase stage of mitosis whereas homologous chromosomes are paired. Standard arrangements of the chromosomes are indicated as +, inverted by capital letters A, D, and E. Probes used for FISH are indicated by the last 5 numbers in red and blue colors in correspondence with the fluorescence dyes Cy3 (red) or Cy5 (blue), which were used for the labeling. Chromosomes are counterstained with YOYO-1 (green). Standard and inverted arrangements are shown for the inversions 1pA (B) and 1qD (D) but standard, inverted, and heterozygous arrangements are shown for the inversion 1qE. In panel D images on the left represent zoomed pictures of the images on the right. Homologous chromosomes are tracked by dashed lines in yellow and red for the standard and inverted arrangements in panel D. The mosquito strains used in this figure are RU3 for +^A^/+^A^, UGA for A/A, RU3 for +^D^/+^D^, OGD for D/D, and MIN for +^E^/+^E^, +^E^/E, and E/E. FISH results provide physical evidence for the presence of the inversions.

In addition, we examined the presence of misassembly 2 and inversions D and E in the 1q arm. To validate the presence of misassembly 2, we performed FISH in 5 individuals from the reference genome strain RU3 (Matthews *et al*. 2018). In all individuals, FISH results showed the reverse order of the AAEL023895 and AAEL025976 probes in the chromosomes compared to the reference genome AaegL5.0, supporting the presence of a misassembly in the reference genome (Fig. S2). Then, we used the same probes to test for overlap between misassembly 2 and inversion 1qD in 2 different strains, UGA and OGD. The Hi-C data mapped to the reference genome indicated the absence of the 1qD inversion in the UGA strain but the presence of this inversion in the OGD strain and the presence of misassembly 2 in the reference genome (Table S3). The FISH results confirmed the presence of the misassembly and absence of the 1qD inversion in the UGA strain. In contrast, FISH results in the OGD strain indicated the presence of only inverted 1qD arrangements in all 3 tested individuals and the presence of misassembly 2 in the reference genome (Fig. 2C).

Finally, we used the same pair of FISH probes, AAEL023895 and AAEL025976, to confirm the presence of 1qE in the PKT and MIN strains. Because inversion 1qE is bigger than inversion 1qD (42 Mb vs 6 Mb), we were able to use proximity to the telomere as an additional marker for interpretation of the FISH results (Fig. 2A). Standard (+^E^/+^E^) and inverted (E/E) arrangements were recognized by signal locations near or distant from the 1q telomere, respectively (Fig. 2D). The heterozygous arrangement (+^E^/E) is shown in the middle panel of Fig. 2D. In this case, signals were observed in different positions on two homologous chromosomes that were paired at the prometaphase stage of mitosis. FISH results in both the PKT and MIN strains showed the presence of standard, heterozygote, and inverted karyotypes of this inversion (Table S3).

Inversions 3pA in the UGA strain and 3pB in the OGD strain were only found in standard and standard/heterozygous arrangements among 4 and 10 tested individuals, respectively (Table S3, Fig. S3). Thus, FISH results provided clear evidence that the inversions identified by the Hi-C method are real.

### Inversion nomenclature and metrics

Because there was no overlap between the inversions we document here and those putatively identified in previous work (Dickson *et al*. 2016; Redmond *et al*. 2020), we established a new nomenclature for the discovered inversions. First, the name of each inversion included the name of the chromosomal arm on which it was found, following the nomenclature for *Ae. aegypti* chromosomes (Sharakhova *et al*. 2011a).

The chromosomal complement in *Ae. aegypti* consists of 3 pairs of chromosomes with chromosomes 1, 2, and 3 being smallest, largest, and intermediate, respectively. All chromosomes are metacentric, with arms almost equal in length. There are no typical heteromorphic X and Y sex chromosomes in *Ae. aegypti*. Instead, sex is determined by a sex locus located on homomorphic chromosome 1, which is morphologically indistinguishable between the sexes (Hall *et al*. 2015; Matthews *et al*. 2018). Chromosomal arms are named as 1p, 1q, 2p, 2q, 3p, and 3q where p is the shorter arm and q is a longer arm. In addition to the chromosomal arm, the name of each inversion includes a capital letter (A, B, C, D, etc.) to avoid confusion with previously described putative micro-inversions that were named by small letters (Redmond *et al*. 2020).

The chromosomal inversions identified in field strains were unequally distributed among chromosomal arms: two inversions (1pA, 1pB) in the 1p arm; five inversions (1qC, 1qD, 1qE, 1qF, and 1qG) in the 1q arm; one inversion (2pA) on arm 2p; three inversions (2qB, 2qC, and 2qD) on arm 2q; six inversions (3pA, 3pB, 3pC, 3pD, 3pE, and 3qF) on arm 3p, and one inversion (3qG) on arm 3q (Fig. 3). Thus, most of the inversions were found on 1q (n=5) and 3p (n=6). Within chromosomal arms, inversions were most dense in the area close to the telomere on arm 1q (regions 1q42–44) and in the middle of arm 3p (regions 3p14–34). The lengths of inversions found in recently colonized strains varied from 5,200,000 bp (3qG) to 54,775,000 bp (3pB) with a mean inversion length of 21,971,389 bp and a median inversion length of 15,985,000 bp (Table 2). The length of two inversions found in the old lab strains were smaller—1,440,000 bp for 2qL in the Liverpool strain and 845,000 bp for 3qR in the RU3 strain, respectively. The inversion found on arm 2p (2pM) in *Ae. mascarensis* has a length of 5,950,000 bp.

**Fig. 3.**
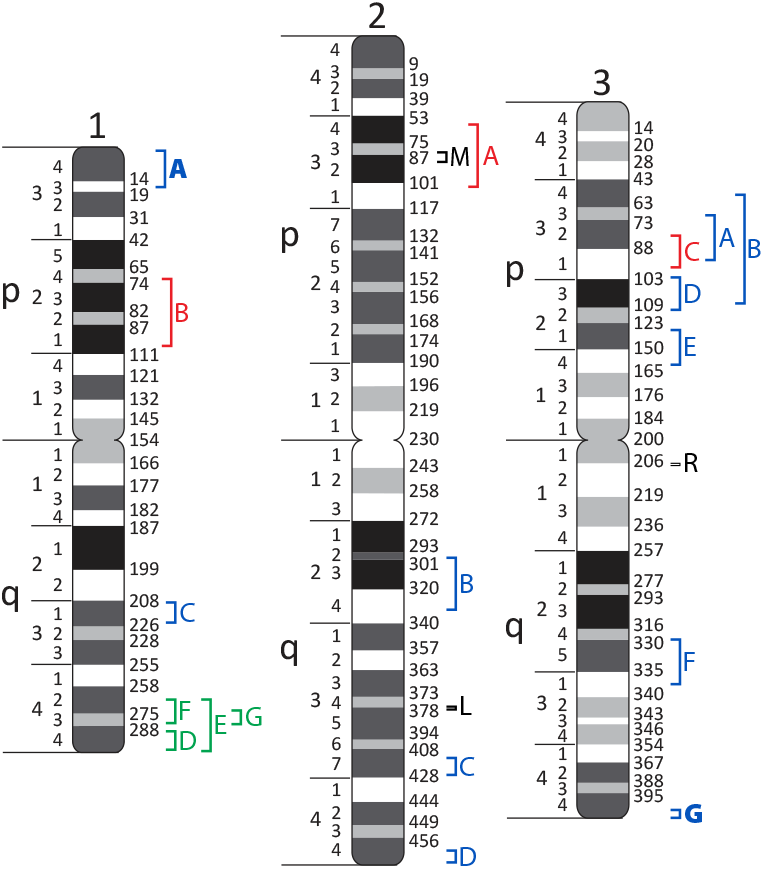
Chromosomal map of the inversion identified in *Aedes aegypti* and *Aedes mascarensis*. Chromosomes are labeled as 1, 2, 3; short and long arms are indicated as p and q, respectively. Numbers on the left indicate chromosome regions and on the right show genome coordinates in Mb. Inversions endemic in West Africa, other inversions in Africa, and inversions found outside of Africa are shown by green, blue, and red colors, respectively. Common African inversions 1pA and 3qG are shown in bold. Inversions 2pM in *Ae. mascarensis*, 3qR and 2qL in RU3 and Liverpool strains are shown black. Inversions are unevenly distributed along the chromosomes whereas most of the inversions localize in the 1q and 3p arms.

### Geographical distribution and population patterns

To better understand the geographical distribution of the chromosomal inversions found in *Ae. aegypti*, we included multiple strains from across the global tropics in this study. Among 23 recently colonized field strains, 12 strains were from 6 countries in Africa: Burkina Faso, Gabon, Kenya, Nigeria, and Uganda, and 11 strains were from 6 countries outside of Africa: Brazil, Colombia, Malaysia, Mexico, Thailand, and the USA. Overall, the study identified more inversions in African strains than in non-African strains: 15 versus 3 inversions, respectively, confirming the ancestral status of African *Ae. aegypti* populations versus non-African (Fig. 4). Among all the inversions found in Africa, 2 inversions, 1pA and 3qG, were abundant in multiple strains distributed across the continent and, therefore, are considered to be common African inversions. Four inversions, 1qD, 1qG, 1qE, and 1qF, were found only in West Africa and were considered to be West African endemic inversions. Another 9 inversions were considered rare because they were found in only one or two African strains. Among 3 chromosomal inversions in *Ae. aegypti* outside of Africa, the 3pC inversion was found in 4 Mexican populations. However, based on the signal outside of the diagonal, in 2 of these populations the frequency of the inversion was low. Intriguingly, inversion 1pB was found in two geographically distant populations in Thailand and in the USA (in Florida), suggesting a potential common ancestry of these strains. Only 1 strain from Africa (SHM in Kenya) and 4 from outside Africa (2 from Florida, USA, 1 from Colombia, and 1 from Malaysia) showed no evidence of any inversions. Thus, inversions in *Ae. aegypti* are abundant and highly associated with the geographical location of the strain.

**Fig. 4.**
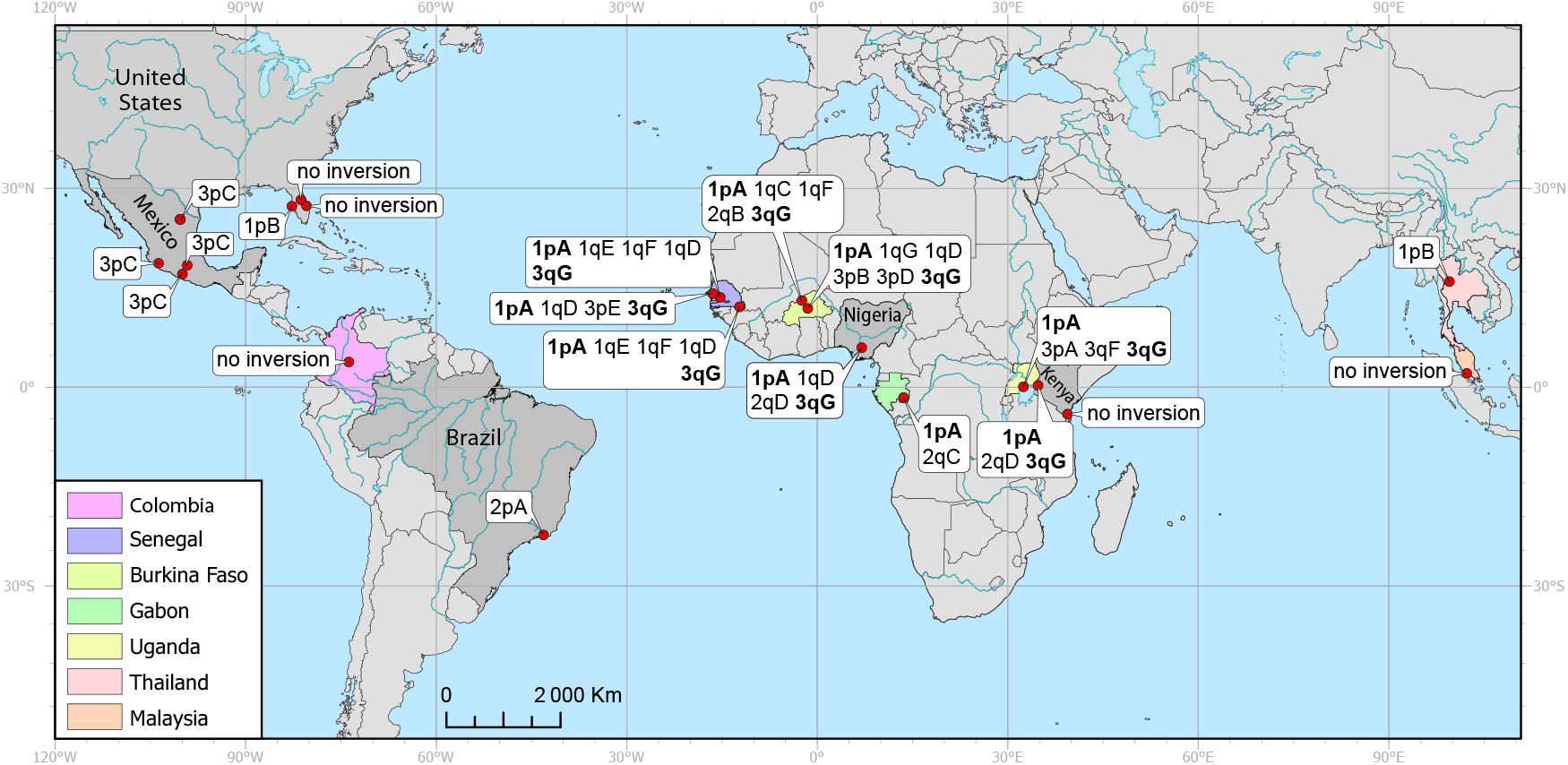
Geographical distribution of the chromosomal inversions discovered by Hi-C analysis. Common inversions 1pA and 3qG in Africa are shown in bold. Countries are shown by different colors. Inversions are more abundant within Africa than outside of Africa. There is no overlap between African and non-African inversions. Inversions were not present in 5 locations.

In addition to characterizing the geographical distribution of each inversion based on Hi-C data from lab colonies, we identified inversion-specific SNPs associated with two cosmopolitan inversions in Africa, 1pA and 3qG. For the SNP identification, we used publicly available whole-genome sequences from field-collected individuals (Rose *et al*. 2020) shown in Table S4. First, to control for the effects of population structure, we examined genetic structure across the two inversion loci within a single geographic region, southern Senegal, which was expected to harbor both inversions at intermediate frequencies. As expected, principal component analyses of inversion loci yielded bi- or tri-modal distributions of individuals across the first principal component, likely corresponding to the different inversion karyotypes (Fig. 5A–B). The SNPs driving this structure showed strong linkage disequilibrium for multiple places across each inversion locus (Fig. 5C–D, Fig. S4), and the consensus genotype across these SNPs for all 393 examined individuals corresponded precisely to the distribution and abundance of inversions found in our Hi-C analyses (Table S4). In particular, we identified putatively diagnostic SNPs with near-perfect correlation to the estimated inversion karyotype (Table 1).

**Fig. 5.**
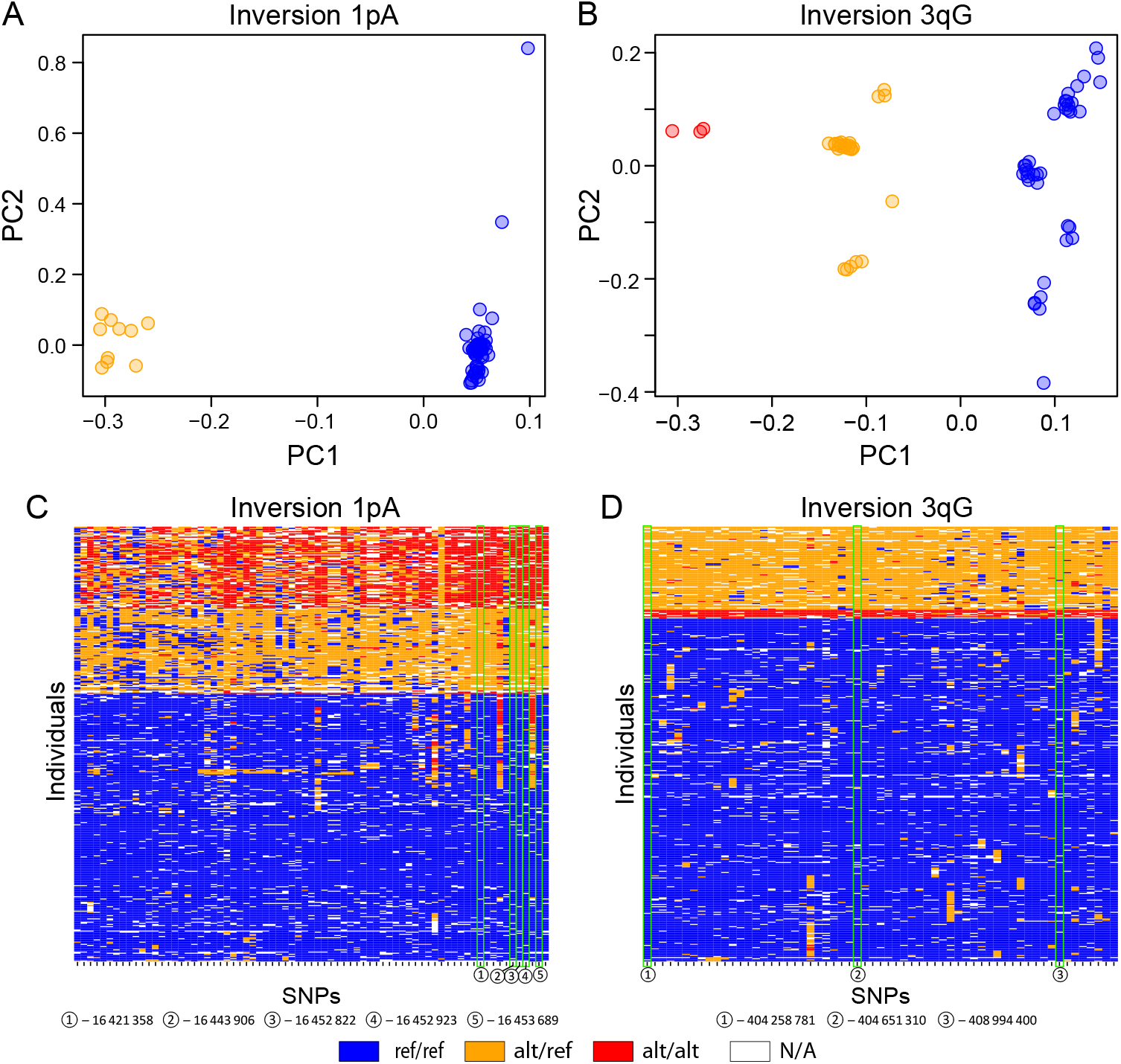
Identification of local genetic structure and inversion-linked SNPs. **A-B**. Principal component analysis (PCA) of biallelic SNPs with minor allele frequency >5% across the inversion 1pA (A) and 1qG (B) loci in southern Senegal identified discrete clusters along PC1. **C-D**. Inversion-linked SNPs. SNPs that were strongly loaded onto PC1 in southern Senegal showed strong linkage disequilibrium across the inversion locus in 393 unrelated individuals, with genotypes corresponding to putative karyotypes. Positions of diagnostic SNPs, for which the individual SNP genotype is in near-perfect agreement (Pearson correlation >0.99) with the consensus genotype across all inversion-linked SNPs are highlighted with green rectangles.

Based on these inversion-linked SNPs, we estimated the frequencies of the standard, inverted, and heterozygote karyotypes for inversions 1pA and 3qG in 31 populations across Africa. The results of this analysis are shown in Fig. 6A. Inversion 1pA was more prevalent in Central Africa while inversion 3qG was present with high frequencies in both Western and Central Africa. We used the chromosomal inversion frequencies of 24 African-wide populations and 2 populations outside of Africa to perform a hierarchical clustering analysis (Fig. S5, Fig. 6B). The analysis subdivided all mosquito populations into four large clusters: 1) Eastern Kenya and non-African locations; Coastal West Africa; 3) West-Africa; and 4) Central Africa. In Africa, most of the populations followed the patterns of their geographical origin with only minor exceptions. The first cluster included 5 populations from Coastal Kenya (KWA, SHM, RAB’, ABK, GND, KBO) and 2 non-African populations from Brazil (SAN), and Thailand (BKK). This cluster was associated with extremely low frequencies or the absence of inversion 1pA and complete absence of inversion 3qG. A population of Central African origin in Gabon (LPV), which also falls into this cluster, had a low frequency of 1pA inversion and no 3qG inversion. The second cluster included 4 populations located in coastal areas of West and Central Africa in Senegal (THI, NGO), Ghana (KUM), and Gabon (LBV). In contrast to East Coastal African populations, 1pA inversion frequencies were relatively high and inversion 3qG was also present. The third cluster included 7 populations located in inland West Africa: Senegal (MIN, PKT, BTT, and KED); Burkina Faso (OHI); Ghana (KIN’ and BOA); and Nigeria (AWK). The presence of both 1pA and 3qG was a characteristic of this cluster. Finally, the fourth cluster included 4 tropical Central African populations from Uganda (BUN’, KCH, ENT), and 2 from Western Kenya (KAK, VMB). A typical feature of this cluster was extremely high frequencies of the 1pA inversion and the presence of the 3qG inversion. One population from Burkina Faso (OGD’) also fell into this cluster because of the abundance of the 1pA inversion in this population. Thus, our study suggests that chromosomal inversions can be used as additional markers to better understand the population structure of *Ae. aegypti* populations.

**Fig. 6.**
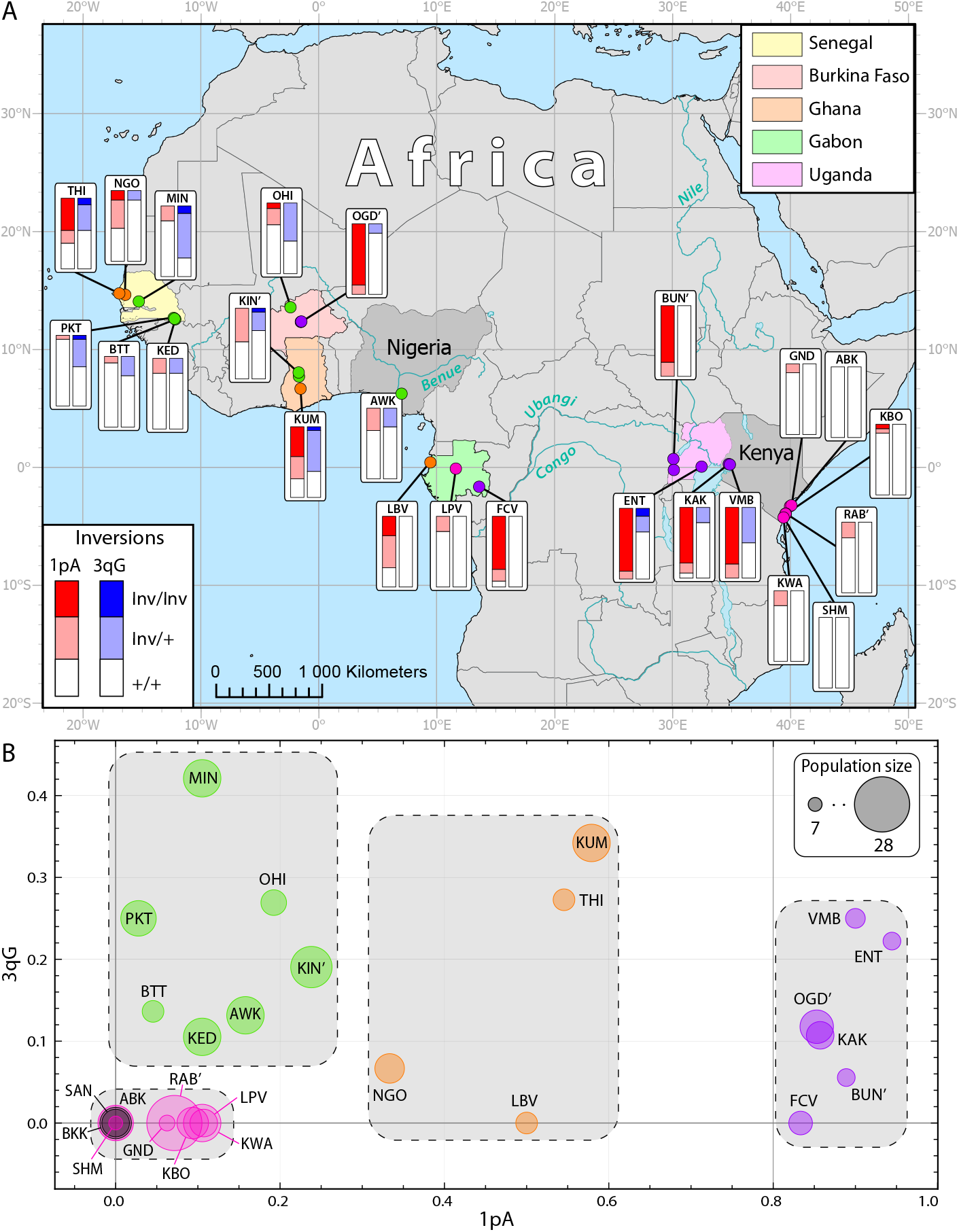
Frequencies of the chromosomal inversions 1pA and 3qG discovered by SNP analysis in whole genome sequences of African mosquitoes. **A**. Frequencies of the inversions in populations across the Africa. Proportions of standard, inverted, and heterozygote arrangements are shown by charts with different colors. Geographical locations are indicated by population ID (Table S4). Colors indicate correspondence of these locations to 4 different clusters shown in panel B. **B**. Clusters associated with inversion frequencies. Populations are shown by circles with different colors and are also indicated by population ID (Table S4). Population sizes are proportional to the diameter of the circles. Colors of these circles indicate correspondence of this location to the 4 clusters (Fig. S5) shown by gray rectangles. In 4 locations, KIN’, OGD’, RAB’, and BUN’, 2 or 3 closely located populations with low number of individuals were combined and named by the population with the biggest number of individuals (Table S4). The composition of clusters correlates with the geographic locations of populations. These clusters correspond to the following geographic regions: 1) coastal East Africa (pink) and non-Africa (gray): 2) coastal West Africa (orange); 3) West Africa (green); 4) Central Africa (purple). Mosquitoes from two locations only, OGD (Burkina Faso) and LPV (Gabon), fall into different clusters than expected in accordance with geographical locations: Central Africa and Coastal East Africa/non-Africa, respectively.

### Potential associations of the inversions with epidemiologically important traits

To explore the potential contribution of chromosomal inversions into epidemiologically important phenotypes in *Ae. aegypti*, such as olfactory behaviors or vector competence, we attempted to find colocalizations of chromosomal inversions, chemoreceptor genes, and QTL related to vector competence. Chemoreceptor genes involved in odor recognition are subdivided into three families: odorant receptors (ORs), gustatory receptors (GRs), and ionotropic receptors (IRs) (Matthews *et al*. 2018). We calculated an overlap significance between each particular inversion and the positions of each class of chemoreceptor genes separately. Based on this computing, we identified 7 inversions that colocalized with chromosomal positions of chemoreceptor genes with sufficient statistical significance (Fig. 7, Table S5). West-African overlapping inversions 1qF and 1qE, and Brazilian inversion 1pA colocalized with the OR genes. Mexican inversion 3pC, and overlapping African inversions 3pA, 3pB, and 3pD colocalized with IR genes. Finally, overlapping African inversion 3pB and 3pD also colocalized with the positions of GR genes. We applied the same calculation to find an overlap between inversions and QTL for various infections (Timoshevskiy *et al*. 2013). The Brazilian inversion 2pA and African inversion 3pB significantly overlapped with QTL related to the transmission of various infectious disease agents by mosquitoes.

**Fig. 7.**
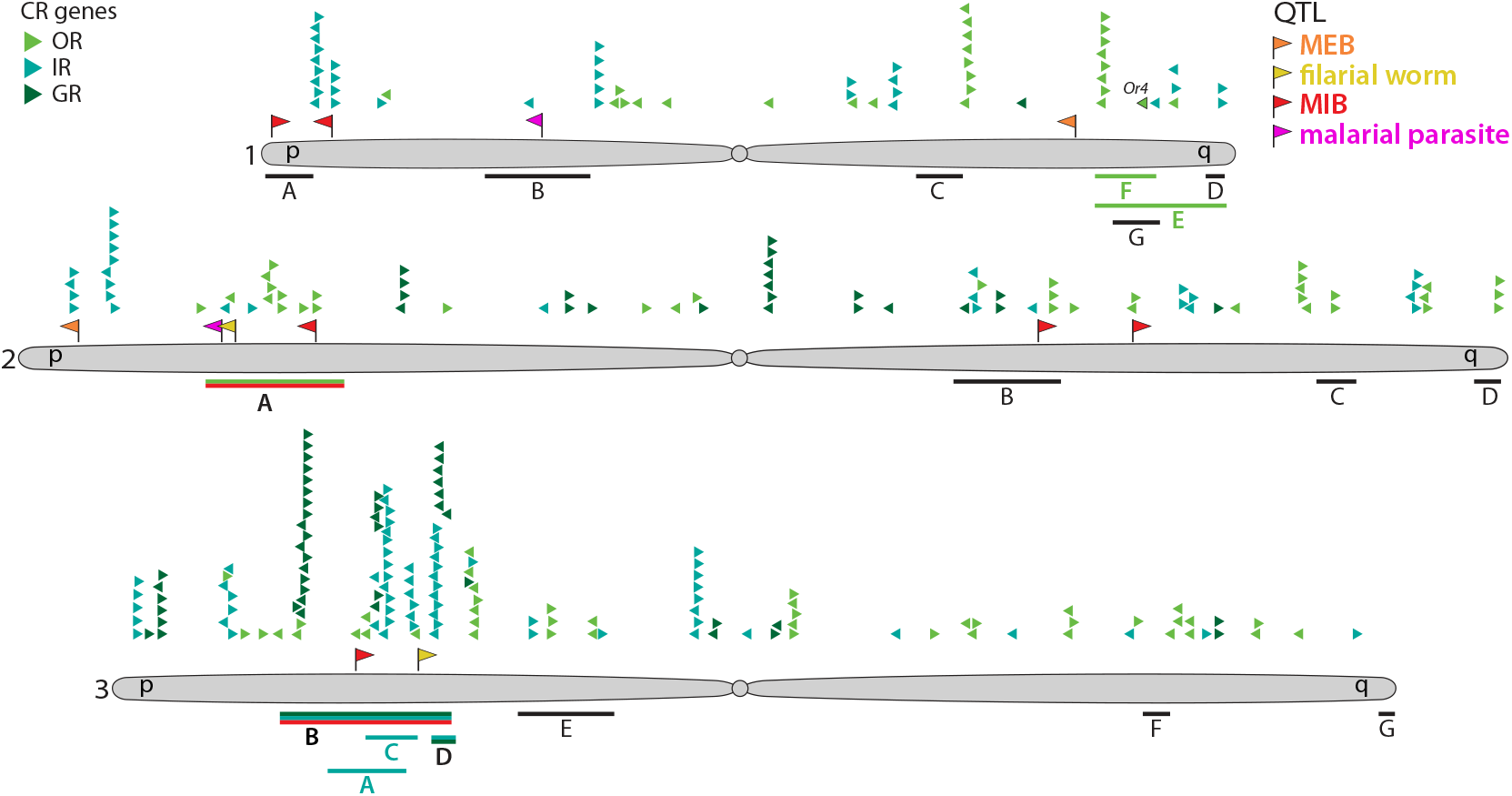
Associations of the chromosomal inversions with chemosensory receptor genes and quantitative trait loci related to vectorial capacity. Chromosomes are indicated by the numbers 1, 2, and 3 and arms as short p and long q. Chemosensory receptors (CRs) are subdivided into odorant receptors, gustatory receptors and ionotropic receptors (ORs, GRs and IRs, respectively). MIB (midgut infection barrier) and MEB (midgut escape barrier) indicate two different types of QTL related to dengue infection. Chromosomal positions of CR genes and QTL are shown by small triangles and flags in different colors, respectively. Chromosomal inversions are shown by colored and black lines below the chromosomes. Inversions associated with CR genes and QTL are labeled with the same colors.

Because chemoreceptors are distributed non-randomly throughout the genome and clustered into chemoreceptor-rich regions, we performed additional calculations to find an overlap between inversions and clusters of evolutionary related chemoreceptors (Ryazansky 2023). We first identified chemoreceptor genes with similar evolutionary trajectories and combined them into evolutionary clusters based on their position on phylogenetic trees and their chromosome positions. For example, 60 out of a total number of 106 OR genes were combined into 24 OR clusters and 46 remaining genes (Table S6). When overlaps between inversions and chemoreceptor gene clusters/ remining genes were calculated, 4 inversions were still identified as enriched with these genes: 1qF with OR genes, 1qC with IR genes, and inversions 3pB and 3pD were enriched with GR genes (Table S7). Finally, we employed a permutation test (Cabrera *et al*. 2012) to validate whether inversion regions were enriched for chemoreceptor genes. We applied 100 circular chromosome permutations, in which a single random offset drawn from a uniform distribution across the length of each chromosome. Circular permutations also indicated a significant overlap (P<0.01) between inversions and chemoreceptor genes taking into account the clustering of chemoreceptor genes within the genome (Fig. S6). Thus, these results suggest potential associations of some of the chromosomal inversions with epidemiologically important phenotypes in *Ae. aegypti* mosquitoes such as olfactory behavior and vector competence.

### Genetic similarities between inversions in *Aedes aegypti* and *Anopheles gambiae*

It has been shown that the 1q and 3p chromosomal arms in *Ae. aegypti* are homologous to the inversion-rich 2R arm in the malaria vector *An. gambiae* (Arensburger *et al*. 2010; Timoshevskiy *et al*. 2014). A total of 7,347 single-copy orthologs were identified between the two species and used to test whether inversions in one species significantly overlap with inversions in the other species (Fig. 8). We compared the location of genes inside of the common inversions of *Ae. gambiae* (Sharakhov *et al*. 2006; Coulibaly *et al*. 2007; George *et al*. 2010; Lobo *et al*. 2010) (Table S8) with locations of the inversions described in this study for *Ae. aegypti*. The results of the hypergeometric test identified 33 significant associations after multiple testing correction out of total of 126 comparisons. The most significant associations were found between the 3pA (UGA) and 3pB (OGD) inversions of *Ae. aegypti* and the 2Rj inversions of *An. gambiae*. Also, the 2pA (B53) inversion of *Ae. aegypti* highly significantly overlapped with the 2La inversion of *An. gambiae*. Moreover, inversion 3pD in *Ae. aegypti* highly significantly overlapped with multiple inversions of 2R chromosomal regions in *An. gambiae* (2Rbk, 2Rc, 2Rd, 2Ru). These results indicate that common sets of genes between these highly diverged species are nonrandomly implicated in the inversion polymorphism and can potentially be important for their adaptation, suggesting a parallel evolution of the two species under similar environmental conditions.

**Fig. 8.**
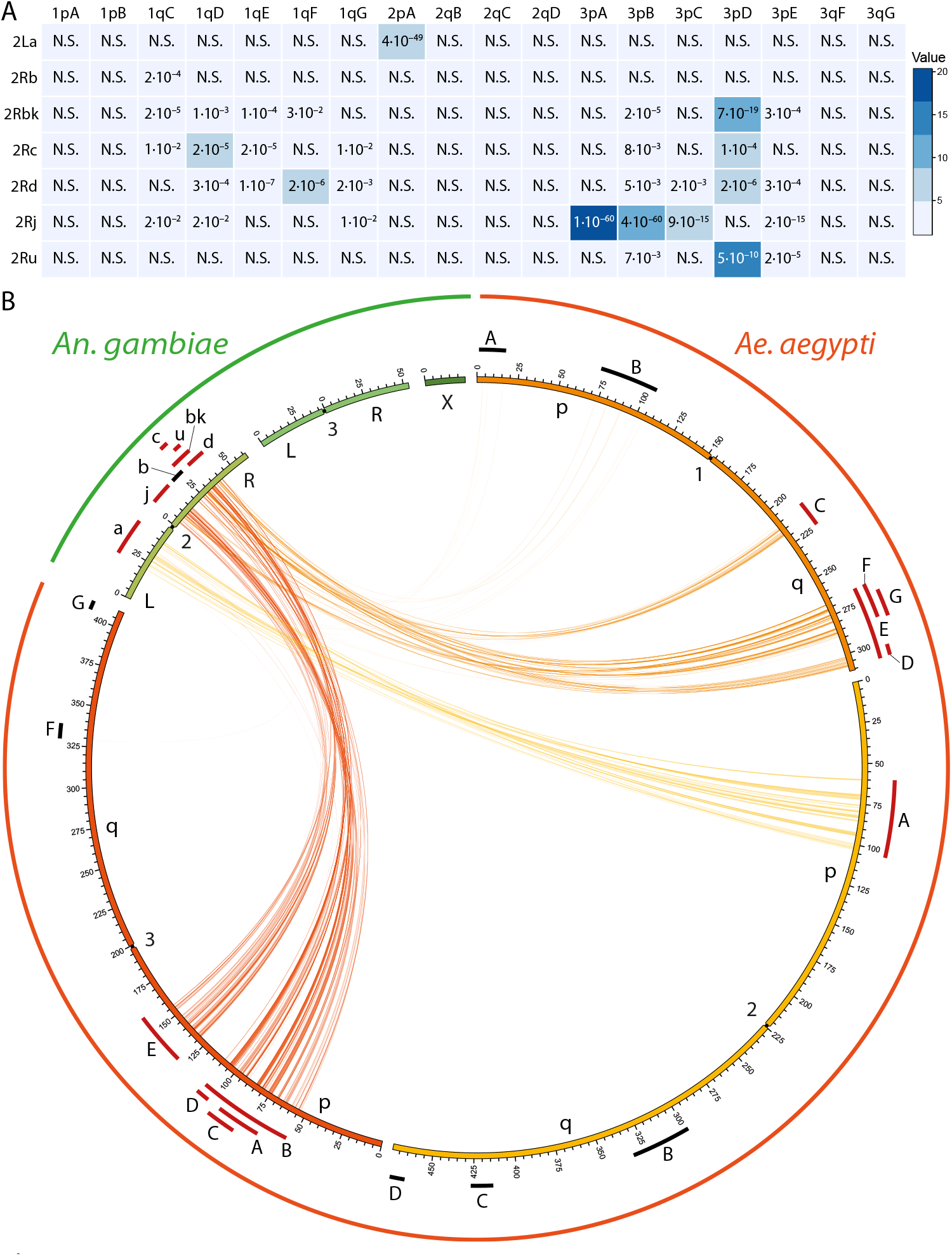
Overlaps between inversions in *Aedes aegypti* and *Anopheles gambiae*. **A**. A matrix of pairwise associations between inversions of *Ae. aegypti* and *An. gambiae* based on the single-copy orthologs. Color intensity and values represent odds ratios and adjusted *P*-values respectively, based on the hypergeometric test. N.S. stands for not significant results. **B**. Circus plot representation of single-copy orthologs between the two species falling into the common regions of inversion polymorphism. Chromosomes are shown by green color for *An. gambiae* and red, orange and yellow colors in *Ae. aegypti*. Chromosomal inversions with significant overlaps between the species are shown by red bars and the rest of the inversions are indicated by black bars. Lines connect positions of the ortholog genes in the genomes of the two species. Inversion-rich regions in the 1p and 3q arms in *Ae. aegypti* significantly overlap with inversions in the 2R arm. Widely spread in Africa, inversion 2La of *An. gambiae* overlaps with inversion 2pA found in *Ae. aegypti* from Brazil.

## Discussion

### Chromosomal inversions in mosquitoes

Although inversion polymorphism in natural populations is well-characterized in malaria-transmitting *Anopheles* mosquitoes, including the major vector of malaria in Africa, *An. gambiae* (Coluzzi *et al*. 2002; Love *et al*. 2016; Love *et al*. 2020), finding chromosomal inversions in *Ae. aegypti* has been challenging due to the difficulty of obtaining high quality polytene chromosome preparations from this species (Campos *et al*. 2003). In this study, for the first time, we document presence of the large chromosomal inversions in the *Ae. aegypti* genome using a molecular method called Hi-C proximity ligation (Corbett-detig *et al*. 2019). All inversions found here are new and do not match previously described putative chromosomal inversions found by FISH (Dickson *et al*. 2016) or micro-inversions discovered using a linked-read sequencing approach (Redmond *et al*. 2020). In fact, the Hi-C approach is not suitable for identification of micro-inversions as their butterfly-like pattern, if present, would be located very close to or “buried” inside the main diagonal on the contact heat map because of the small inversion size under 500 kb. Interestingly, the development of a linkage map based on hybrids between the *Ae. aegypti* Liverpool strain and the PK-10 strain of *Ae. aegypti* from Senegal indicated severely reduced recombination in F2 offspring, suggesting the presence of four, one, and three inversions in chromosomes 1, 2, and 3, respectively (Bernhardt *et al*. 2009). In our study, we discovered four inversions in chromosome 1 and one inversion in chromosome 3 in the PKT strain collected from the same area as the PK-10 strain in Senegal (Rose *et al*. 2020). One and two inversions were found in chromosomes 2 and 3, respectively, in other strains from West Africa. Although the genomic positions of the genetic markers used in the linkage mapping study did not match with the inversion positions found here, it is likely that the inversions contribute to reduced recombination in the chromosomes in hybrids between the Liverpool and PK-10 strains.

Our study identified a total of 20 inversions in the *Ae. aegypti* genome: 18 in recently colonized strains and two in old laboratory strains. An additional inversion was found in the closely related species *Ae. mascarensis*. The lengths of chromosomal inversions in natural populations of *Ae. aegypti* varied between 5 and 55 Mb, nearly twice the length of those in *An. gambiae* (4–22 Mb) (Love *et al*. 2019). Such differences are likely related to the fact that the *Ae. aegypti* genome is 4.4 times the size of *An. gambiae*, 1.25 Gb (Matthews *et al*. 2018) versus 0.25 Gb (Holt *et al*. 2002), respectively. As in the *An. gambiae* genome, inversions in *Ae. aegypti* were unevenly distributed along chromosomal arms. In *Ae. aegypti* the highest numbers of inversions were identified in the 1q and 3p arms. In *An. gambiae*, most chromosomal inversions have been found in arm 2R, which is homologous to both the 1q and 3p arms of *Ae. aegypti* (Timoshevskiy *et al*. 2014).

Based on the frequency of each inversion in natural populations, we classified them into three categories: common (present in multiple geographically distinct populations), endemic (restricted to a particular geographic region), and rare (found in only one or two populations). Among 15 inversions found in Africa, only two (1pA and 3qG) were common across the entire continent. Interestingly, only two *An. gambiae* inversions, 2La and 2Rb, are widely spread in Africa (Love *et al*. 2019). Most of the endemic chromosomal inversions in *Ae. aegypti* (1qF, 1qD, 1qE, and 1qG) were found in West Africa. Similarly, three chromosomal inversions (2Rc, d, and u) on the 2R arm of *An. gambiae*, which is homologous to 1q arm of *Ae. aegypti*, are restricted to West Africa and only rarely found in Central and Eastern Africa. However, these species comparisons are tentative as much better information on geographical distribution of inversions in *Ae. aegypti* is needed to reach any strong conclusions.

Nevertheless, it is tempting to speculate that the similar geographic distribution of the inversion polymorphism in two distant African mosquitoes, *Ae. aegypti* and *An. gambiae*, is related to their shared natural environment. If the same genes are captured by inversions in both species, they may have the same phenotypic effects. In this study, we directly examined overlap between the genes located inside *Ae. aegypti* and *An. gambiae* inversions. Interestingly, multiple genes inside of the endemic West African *An. gambiae* inversions 2Rc, 2Rd, 2Ru (Love *et al*. 2019) overlapped significantly with the 3pD inversion of *Ae. aegypti*, which is also endemic to West Africa (found in OGD, Fig. 8). Also, significant associations were found between the 3pA and 3pB inversions of *Ae. aegypti* and the 2Rj inversion of *An. gambiae* (Fig. 8). Although we did not find correlations between the common inversions 1pA and 3qG in *Ae. aegypti* and the 2La and 2Rb in *An. gambiae* in Africa, we found a strong association between the common African 2La inversion in *An. gambiae* and the 2pA inversion in *Ae. aegypti* in Brazil. Overall, these results indicate that some inversions segregating in these distantly related mosquito vectors nonrandomly capture common sets of genes that may be important for mosquito adaptation, suggesting a parallel evolution of the two species under similar environmental conditions. Alternatively, these genomic regions may share properties that make them unstable leading to a higher-than-average rate of production of chromosomal rearrangements. A comparative study of chromosomal inversions in *An. gambiae, An. funestus*, and *An. stephensi* indicated that all three species have the highest number of inversions on the 2R arm and capture similar sets of genes, although the break-points were different (Sharakhova *et al*. 2011b). A recent review highlighted involvement of chromosomal inversions in parallel evolution in different organisms (Westram *et al*. 2022). Such parallel evolution is most likely to be detected in regions subjected to strong selection such as inversions.

Interestingly, in two populations of *Ae. aegypti* in Senegal, we found an unusual rearrangement, a triple inversion on arm 1q comprising three overlapping inversions 1qD, 1qF, and 1qE. Although the 1qD and 1qF inversions were sometimes found alone, inversion 1qE always included the other two, indicating that the 1qE inversion originally occurred on a chromosome carrying both of the other inversions. Interestingly, in the malaria mosquito *An. messeae* a similar rearrangement, inversion 3L1, is represented by a combination of at least two inversions (Artemov *et al*. 2021). This double inversion is widespread in natural populations in Eurasia but the two single inversions inside 3L1 were never described separately in nature (Brusentsov *et al*. 2023). Such overlapping chromosomal inversions should lead to a greater reduction of recombination compared to single inversions. Indeed, the triple inversion 1qE overlaps with the highest peaks of genetic divergence between strains in West Africa (Rose *et al*. 2020). Such suppression of recombination may cause differentiation of the gene arrangements and may accelerate the accumulation of fixed genetic differences among populations. Thus, our discovery of chromosomal inversions in *Ae. aegypti* will assist in adequate interpretation of genomic data in this species.

### Inversions and population genetics of *Aedes aegypti*

According to recent investigations, *Ae. aegypti* likely originated in the islands of the South West Indian Ocean after diverging from its closest relative *Ae. mascarensis* ≈7 MYA and invading Africa ≈85,000 years ago (Soghigian *et al*. 2020). Beginning about 5,000 years ago in West Africa, two forms began to emerge, one remaining in its native forest ecotone habitat and the other beginning to use human settlements to survive long dry periods (Rose *et al*. 2023). This latter form was pre-adapted to life aboard ships and began to spread around the tropical and sub-tropical regions in the world about 500 years ago. Gradually, this human-associated form acquired the morphological and behavioral traits we associate with subspecies *Aaa* (Mattingly 1957; Brown *et al*. 2011; Brown *et al*. 2014; McBride *et al*. 2014; Gloria-soria *et al*. 2016; Powell *et al*. 2018; Rose *et al*. 2020).

Our study revealed a striking difference between African inversion-rich populations and non-African inversion-poor populations supporting a basal status of African populations of *Ae. aegypti* versus non-African populations. Although the total number of strains examined was almost the same for African and non-African populations (12 versus 11, respectively), the majority of inversions were found in African strains (15 versus 3). Interestingly, there was no overlap between inversions found in African and non-African populations. This could be due to insufficient sampling in our study or, perhaps, new inversions arose or drifted to high frequency during population bottlenecks in *Aaa* over the relatively short period (≈500 years) since it escaped Africa. Thus, the inversion data add to the growing support for the genetic distinctness of *Aaa* and *Aaf*, with *Aaf* being ancestral and *Aaa* being monophyletic, implying a single out of Africa event (Brown *et al*. 2014; Gloria-soria *et al*. 2016; Kotsakiozi *et al*. 2018; Powell *et al*. 2018).

Could inversions provide insights on the geographic origin of the introduction? The ancestral proto-*Aaa* form should have a similar chromosomal arrangement to non-African populations with a low number of inversions or no inversions in their genome. While there are many endemic inversions segregating in West Africa, they have low frequency in some strains in northern Senegal where proto-*Aaa* has been found. For example, many mosquitoes from the human-preferring NGO population have no inversions. Our data are, thus, consistent with the idea that proto-*Aaa* arose in West Africa and was then perhaps introduced to other coastal African slave trading ports, such as Angola, where they mixed further with local populations before migrating to the Americas. Coastal Angola as a place of proximate origin of the non-African *Aaa* has been proposed partly due to the large amount of slave-trade from what is today coastal Angola (Kotsakiozi *et al*. 2018; Rose *et al*. 2023).

In addition, we tested whether geographical clusters associated with the two most common inversions in Africa, 1pA and 3qG, followed the patterns of population structure in *Ae. aegypti* or whether these clusters have their own geographical patterns, suggesting involvement of these inversions in climatic or ecological adaptations of the mosquitoes. Frequencies of the two common inversions were estimated in 31 populations across Africa using the SNP-based genotyping approach developed in this study. The study revealed the presence of 4 major clusters: 1) Eastern Kenya and non-African locations; 2) Coastal West Africa; 3) West-Africa; and 4) Central Africa. In general, these data overlap with the previous observation about the presence of three major geographical clusters in Africa associated with Western, Eastern, and Central Africa (Rose *et al*. 2020). The presence of an additional coastal cluster in West Africa may suggest involvement of these inversions in climate adaptation. Genotype data across thousands of SNPs can provide strong inferences of population structure and historical relationships between populations or species.

Most analytical methods to reach these inferences assume little or no selection on the SNP variants, an assumption that, if violated only infrequently across thousands of SNPs, should not greatly affect results. However, if inversions subject to strong selection are frequent in the genome of a species, the otherwise neutral SNPs captured between breakpoints would be indirectly affected by selection and, thus, violate assumptions of neutrality. Better sampling is required to determine the impact of inversions on neutrality assumptions made for population genetics in *Ae. aegypti*.

### Inversions and behavior

To evaluate how inversions may be affecting traits important in disease transmission, we attempted to correlate locations of the chromosomal inversions with the positions of clusters of chemoreceptor genes involved in smell and taste (Matthews *et al*. 2018) and QTL related to levels of pathogen infections (Bosio *et al*. 2000; Severson *et al*. 2002; Timoshevskiy *et al*. 2013). We found significant overlap of the 1qF inversion, which is endemic in West Africa, with clusters of odorant receptors located close to the telomere on 1q (Matthews *et al*. 2018). As was shown before, this region overlaps with the highest peaks of SNP differences between strains in West Africa (Rose *et al*. 2020), which includes an odorant receptor gene (Or4) that was previously linked to preference for human odors (McBride *et al*. 2014). In addition, the overlapping inversions 3pB/3pD and the inversion 3pC located in the middle of the 3p arm overlapped with the clusters of gustatory and ionotropic receptor genes, respectively. Finally, inversions 2pA and 3pB colocalized with the location of QTL implicated to affect multiple pathogen infections including dengue, filariasis, and, in the case of 2pA, avian malaria (Timoshevskiy *et al*. 2013). Interestingly, inversion 2pA is in a homologous position to inversion 2La in *An. gambiae*, which is known to be associated with malaria transmission (Riehle *et al*. 2017). These results suggest that chromosomal inversions can potentially regulate the behavior of mosquitoes related to their ability to recognize humans as potential hosts and to transmit epidemiologically important pathogens.

## Conclusions

Using Hi-C proximity ligation, we identified 21 multi-megabase inversions in 23 strains of *Ae. aegypti* from across the species’ global distribution, two old laboratory colonies, and the closely related species *Ae. mascarensis*. All inversions discovered in this study were new; thus, we created a new nomenclature for chromosomal inversions in *Ae. aegypti*. The lengths of inversions detected in recently colonized strains varied from 5 to 55 Mbp with a mean of 22 Mbp. Strikingly, three overlapping inversions were found in the 1q arm of 2 strains from West Africa.

Inversions were unevenly distributed along chromosomal arms with the highest numbers determined in the 1q and 3p arms, which are homologous to the inversion-rich 2R chromosomal arm in *An. gambiae*. Direct comparison of inversions between *An. gambiae* and *Ae. aegypti* revealed significant overlap in their genomic positions. This result could account for the parallel evolution of the two species under similar environmental conditions. All chromosomal inversions, including the one specific for *Ae. mascarensis*, were polymorphic. The inversions were highly associated with the geographical origin of the strains and were more abundant in African than in non-African strains, 15 versus 3 inversions, respectively. In Africa, the highest number of inversions was observed in western areas of the continent. The hierarchical cluster analysis, based on 2 highly polymorphic inversions, 1pA and 3qG, subdivided all mosquito populations into four large clusters: 1) Eastern Kenya and non-African locations; 2) Coastal West Africa; West-Africa; and 4) Central Africa suggesting involvement of these inversions in climatic adaptations in Africa. Three inversions in 1p and 3p arms were associated with genomic locations of chemoreceptor genes involved in mosquito behavior. One large chromosomal inversion in the 2p arm overlapped with QTL asociated with susceptibility to dengue virus and filarial worm infections. Thus, our study revealed a large pool of structural variation in the *Ae. aegypti* genome potentially involved in mosquito adaptation to humans and interactions with pathogens.

## Data availability

The raw sequencing data for all Hi-C libraries have been deposited into the NCBI SRA database with accession number: PRJNA1003935. All assembled Hi-C heatmap files are accessible through the GEO series accession number GSE243024 https://www.ncbi.nlm.nih.gov/geo/query/acc.cgi?acc=GSE243024.

## Supplementary figures

Data not shown

**Fig. S1. | Visualization of all inversions detected on Hi-C contact maps in *Aedes aegypti* and *Aedes mascarensis* strains**. The Hi-C maps for the following inversions are shown: the 1pA inversion in the UGA strain (A); the 1pB inversion in the T51 strain (B); the 1qC inversion in the OHI strain (C); the 1qD inversion in the OGD strain (D); the 1qE inversion in the PKT strain (E); the 1qF inversion in the OHI strain (F); the 1qG inversion in the OGD strain (G); the 2pA inversion in the B53 strain (H); the 2qC inversion in the OHI strain (I); the 2qE inversion in the FCV strain (J); the 2qF inversion in the AWK strain (K); the 3pA inversion in the UGA strain (L); the 3pB inversion in the OGD strain (M); the 3pC inversion in the GUER strain (N); the 3pD inversion in the OGD strain (O); the 3pE inversion in the NGO strain (P); the 3qF inversion in the UGA strain (Q); the 3qG inversion in the UGA strain (R); the 2qL inversion in the LVP strain (S); the 3pR inversion in the RU3 strain (T); and the 2pM inversion in the MASC strain of *Ae. mascarensis* (U). Black arrows point to the center of butterfly-like patterns showing the break point location. The color range scheme is shown at the bottom right corner. The enlarged image on the right/left corresponds to the zoomed-in inversion fragment of the main heat map.

**Fig. S2.**
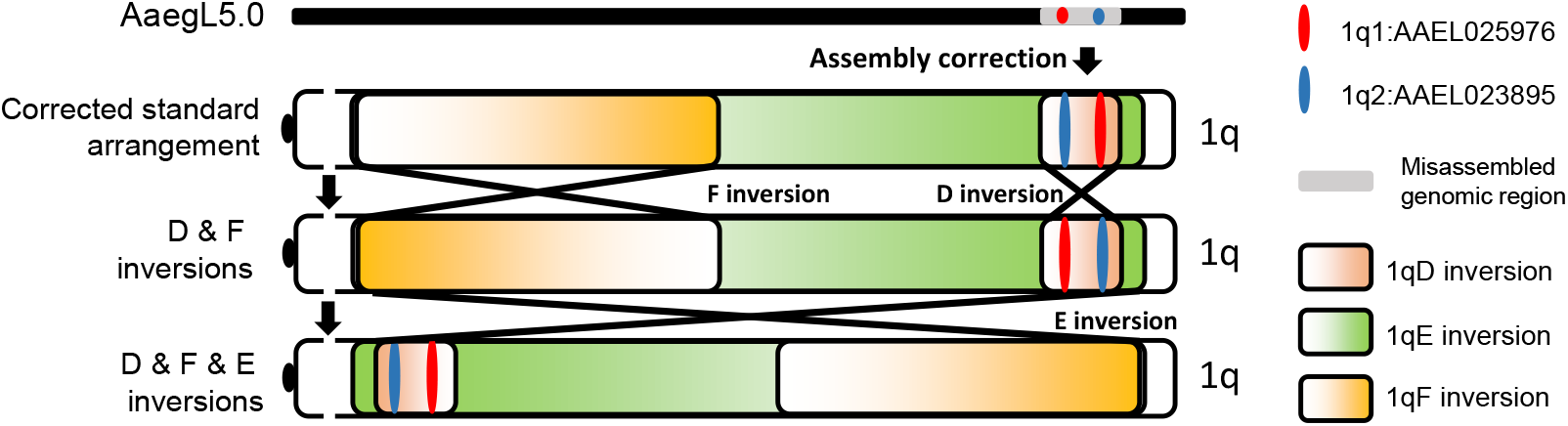
A scheme for correction of misassembly 2 in the 1p arm in the *Ae. aegypti* genome. The corrected order of two genes AAEL023895 (blue) and AAEL025976 (red), which have been used in FISH experiments in this region, are shown within standard and inverted chromosomes.

**Fig. S3.**
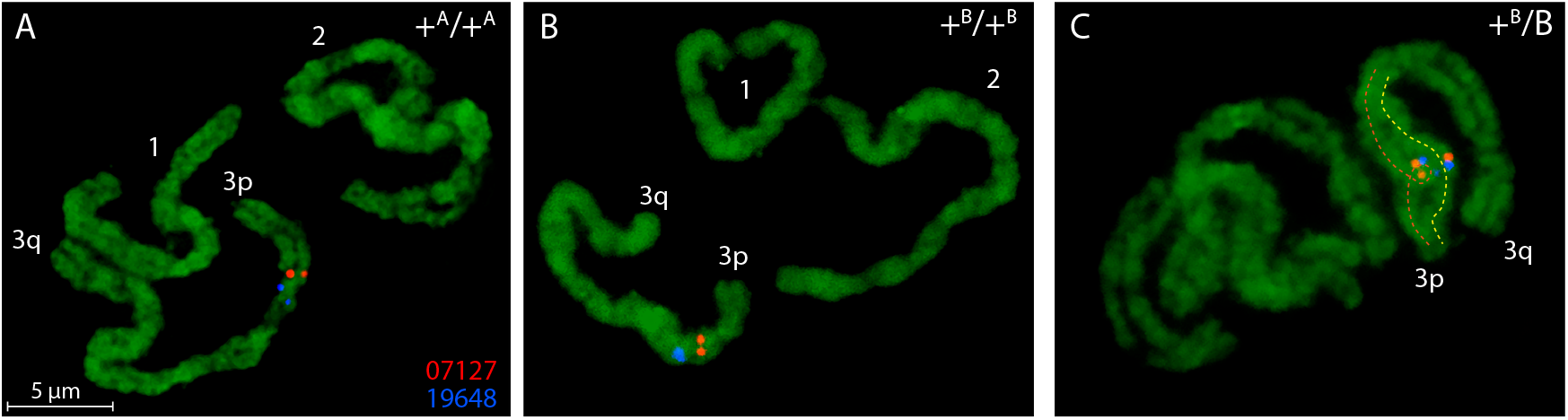
Examples of FISH experiments for detection of the 3pA and 3pB inversions. **A**. Standard arrangement of 3pA in the UGA strain. **B**. Standard arrangement of 3pB in the OGD strain. **C**. A heterokaryotype of the 3pB inversion in the OGD strain. **D**. An inverted grayscale image of the chromosome in figure C. The homologous chromosomes in the inversion loop are tracked with dashed lines of different colors. AAEL007127 (shortened to 07127) is labeled with Cy3 (red), AAEL019648 (shortened to 19648) is labeled with Cy5 (blue).

**Fig. S4.**
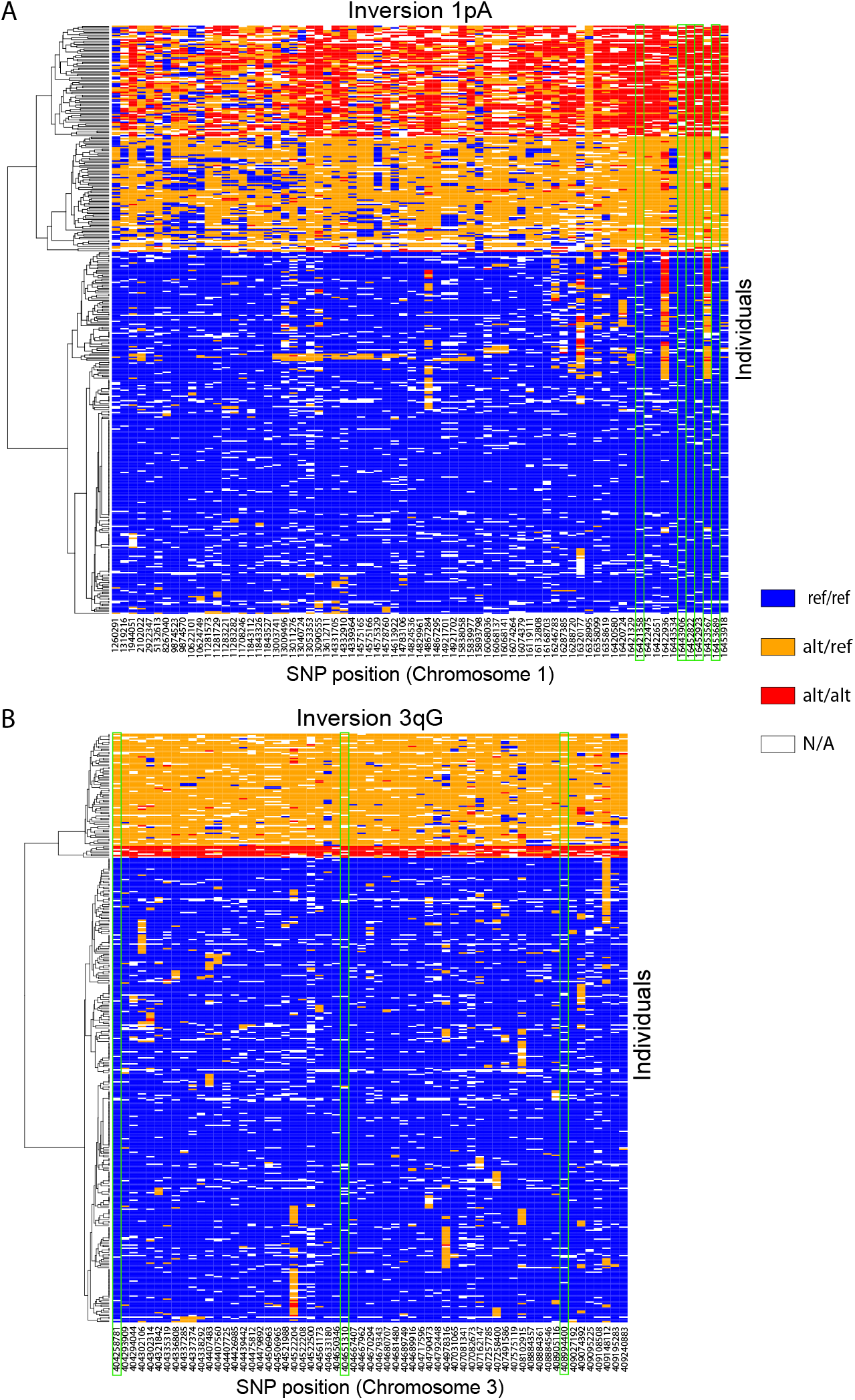
Identification of diagnostic SNPs for the 1pA (A) and 3qG inversions (B). Across 393 unrelated individuals, SNPs that were strongly loaded onto PC1 across the 1pA locus among samples from southern Senegal showed strong linkage disequilibrium across the locus, with genotypes corresponding to putative karyotypes. Positions of diagnostic SNPs, for which the individual SNP genotype is in near-perfect agreement (Pearson correlation >0.99) with the consensus genotype across all inversion-linked SNPs are highlighted with green rectangles. SNP loci are hierarchically clustered by Euclidian distance using complete linkage clustering, and SNPs are displayed as columns across the locus with coordinates indicated in bp. Diagnostic SNPs are indicated by green rectangles.

**Fig. S5.**
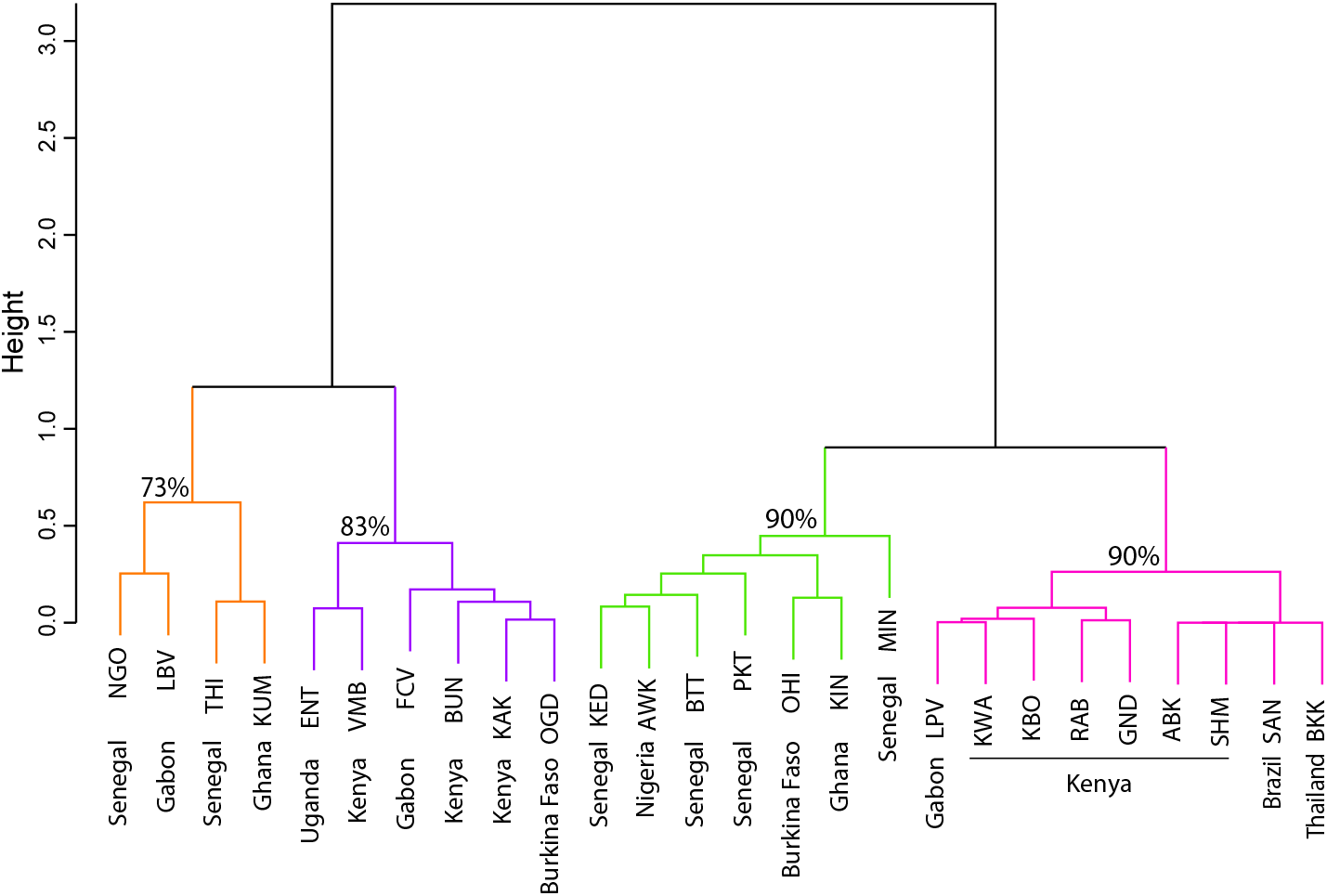
Hierarchical cluster analysis based on frequencies of the 1pA and 3qG inversions in African populations. Four clusters are shown using different colors according to their geographical location: 1) Coastal East African, non-African; 2) Coastal West African; 3) West African; 4) Central Africa. Approximate unbiased P-values are shown for the clusters (Shimodaira 2004). Only the OGD population from Burkina Faso and the LPV population from Gabon did not match their expected geographical regions.

**Fig. S6.**
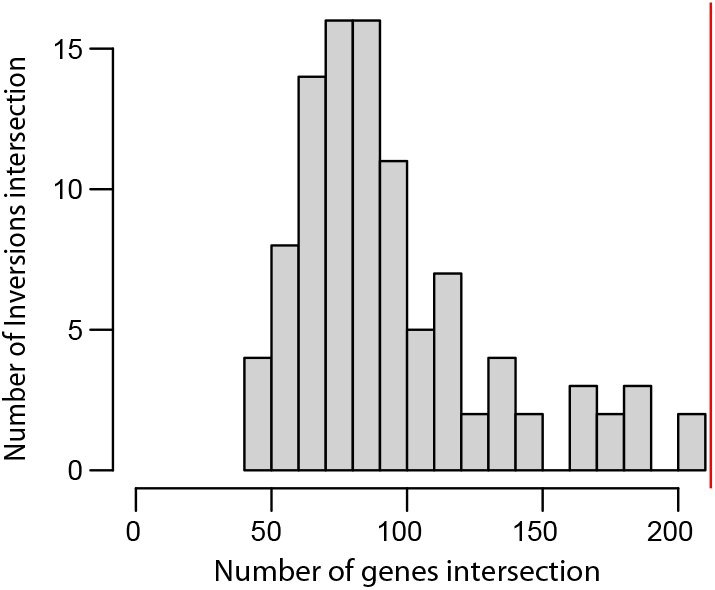
Histogram of the number of chemoreceptor-inversion overlaps across circular chromosomal permutations. The red line showes the true observed number of intersections.

## Supplementary tables

**Table S1.** Mosquito samples used for the Hi-C library preparation. Latitude and longitude are the average coordinates of collection sites in one region. N/A means not applicable or missing information.

| Strain ID | Species | Collection location | Country | Latitude | Longitude | Generations for Hi-C | Numbers of mosquitoes used for Hi-C | Strain information |
| --- | --- | --- | --- | --- | --- | --- | --- | --- |
| NGO | *Aedes aegypti* | Ngoye | Senegal | 14.63960875 | -16.43174115 | g4 | 120 males | Rose et al. 2020 |
| MIN | *Aedes aegypti* | Mindin | Senegal | 14.07473585 | -15.28339615 | g4 | 120 males | Rose et al. 2020 |
| PKT | *Aedes aegypti* | PK10 | Senegal | 12.60970505 | -12.2474927 | g4 | 120 males | Rose et al. 2020 |
| OGD | *Aedes aegypti* | Ouaga-dougou | Burkina Faso | 12.37455556 | -1.49805556 | g4 | 120 males | Rose et al. 2020 |
| OHI | *Aedes aegypti* | Ouahigouya | Burkina Faso | 13.583331 | -2.416665 | g4 | 120 males | Rose et al. 2020 |
| AWK | *Aedes aegypti* | Awka | Nigeria | 6.2509878 | 7.00215995 | g4 | 120 males | Rose et al. 2020 |
| FCV | *Aedes aegypti* | Franceville | Gabon | -1.63997525 | 13.582982 | g7 | 120 males | Rose et al. 2020 |
| UGA | *Aedes aegypti* | N/A | Uganda | N/A | N/A | N/A | 120 males | N/A |
| KAK | *Aedes aegypti* | Kakamega | Kenya | 0.2845454 | 34.7543336 | g6 | 120 males | Rose et al. 2020 |
| SHM | *Aedes aegypti* | Shimba | Kenya | -4.25009085 | 39.4006515 | g6 | 120 males | Rose et al. 2020 |
| K2 | *Aedes aegypti* | Rabai | Kenya | -3.93661547 | 39.57475039 | N/A | >1000embryos | McBride et al. 2014 |
| K27 | *Aedes aegypti* | Rabai | Kenya | -3.924119 | 39.591141 | N/A | >1000embryos | McBride et al. 2014 |
| MONT | *Aedes aegypti* | Monterrey | Mexico | N/A | N/A | N/A | >1000embryos | Garcia-Luna et al. 2018 |
| GUER | *Aedes aegypti* | Guerrero | Mexico | N/A | N/A | N/A | >1000embryos | Garcia-Luna et al. 2018 |
| CUER | *Aedes aegypti* | Cuernavaca Morelos | Mexico | 18.939599 | -99.198412 | g0 | 60 adults | This study |
| COLI | *Aedes aegypti* | Colima Colima | Mexico | 19.24333 | -103.72472 | g0 | 26 adults | This study |
| F54 | *Aedes aegypti* | Anna Maria Island, Florida | USA | 27.52 | -82.72 | g0 | 120 males | Metz et al. 2023 |
| VEBE | *Aedes aegypti* | Vero Beach | USA | 40.738047 | -73.996341 | g0 | 109 males | This study |
| ORL | *Aedes aegypti* | Orlando | USA | N/A | N/A | N/A | >1000 embryos | N/A |
| C62 | *Aedes aegypti* | Acacias, Meta | Columbia | 3.99 | -73.766 | g11 | 120 males | Metz et al. 2023 |
| B53 | *Aedes aegypti* | Paqueta Island | Brazil | -22.759 | -43.109 | g14 | 120 males | This study |
| T51 | *Aedes aegypti* | Muang District, Kamphaeng Phet | Thailand | 16.5 | 99.6 | g13 | 120 males | Rose et al. 2020, Metz et al. 2023 |
| M69 | *Aedes aegypti* | Melaka | Malaysia | 2.24 | 102.26 | g3 | 120 males | This study |
| LVP | *Aedes aegypti* | Lab strain Liverpool | West Africa | N/A | N/A | N/A | 120 males | Severson et al. 2002 |
| RU3 | *Aedes aegypti* | Lab strain Liverpool | N/A | N/A | N/A | N/A | >1000embryos | Matthews et al. 2018 |
| MASC | *Aedes mascarensis* | Le Dauge, Port-Louis | Mauritius | N/A | N/A | N/A | 120 males | N/A |

**Table S2.**
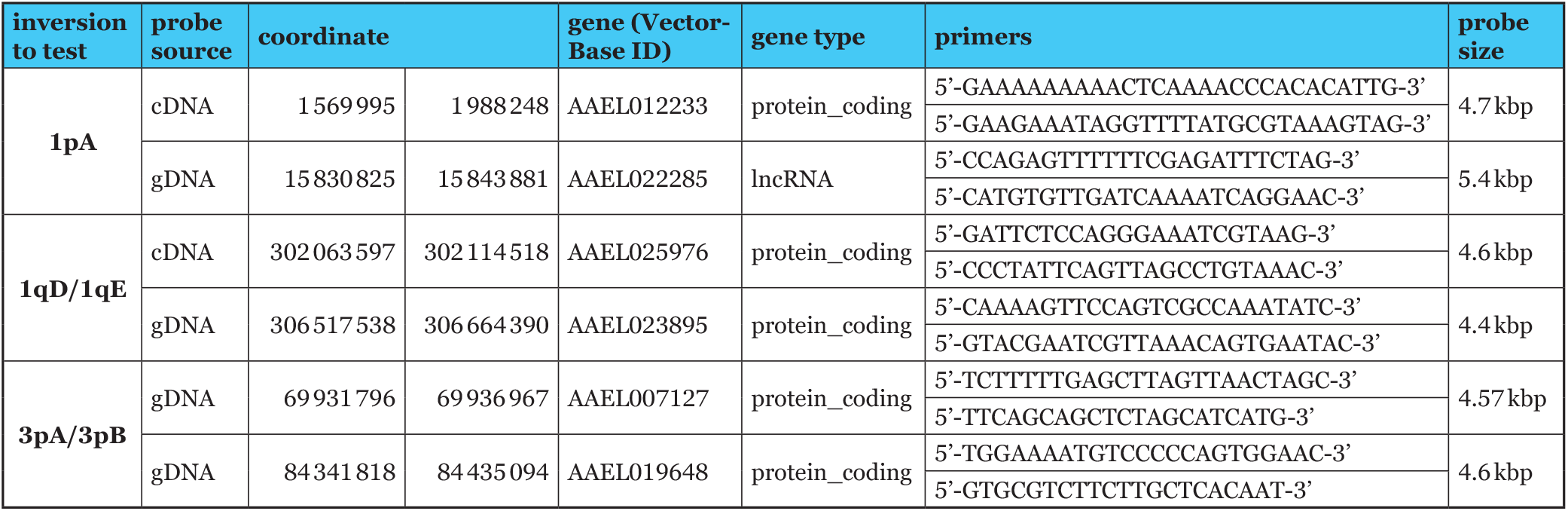
Probes for validation of the chromosomal arrangements using FISH.

**Table S3.** Summary of FISH results in selected mosquito strains. + indicates a standard arrangement of the corresponding inversion.

| **FISH result summary using probe AEEL012233 and AEEL022285** |  |  |  |  |
| --- | --- | --- | --- | --- |
| **Strain ID** | **1pA** | **1pA/+** | **+** | **Total** |
| **RU3** | 0 | 0 | 3 | 3 |
| **PKT** | 0 | 0 | 3 | 3 |
| **UGA** | 7 | 0 | 0 | 7 |

| **FISH result summary using probe AEEL025976 and AEEL023895** |  |  |  |  |
| --- | --- | --- | --- | --- |
| **Strain ID** | **1qD** | **1qD/+** | **+** | **Total** |
| **RU3** | 0 | 0 | 5 | 5 |
| **UGA** | 0 | 0 | 7 | 7 |
| **OGD** | 3 | 0 | 0 | 3 |
|  | **1qE** | **1qE/+** | **+** | **Total** |
| **PKT** | 2 | 2 | 2 | 6 |
| **MIN** | 2 | 3 | 1 | 6 |

| **FISH result summary using probe AEEL007127 and AEEL019648** |  |  |  |  |
| --- | --- | --- | --- | --- |
| **Strain ID** | **3pA** | **3pA/+** | **+** | **Total** |
| **UGA** | 0 | 0 | 4 | 4 |
|  | **3pB** | **3pB/+** | **+** | **Total** |
| **OGD** | 0 | 1 | 9 | 10 |

**Table S4.** Frequencies of the inversions 1pA and 3qG, and Hardy-Weinberg equilibrium (HWE) in the mosquito populations. Populations with a significant deviation from the HWE (P<0.05) are absent.

| Popula-<br>tion ID | Merged popula-<br>tion | 1pA |  |  | 3qG |  |  | HWE P-value |  |
| --- | --- | --- | --- | --- | --- | --- | --- | --- | --- |
|  |  | AA | AB | BB | AA | AB | BB | 1pA | 3qG |
| ABK |  | 18 | 0 | 0 | 18 | 0 | 0 | 1 | 1 |
| AWK |  | 13 | 6 | 0 | 14 | 5 | 0 | 1 | 1 |
| BKK |  | 16 | 0 | 0 | 16 | 0 | 0 | 1 | 1 |
| BTT |  | 10 | 1 | 0 | 8 | 3 | 0 | 1 | 1 |
| BUN' | BUN+KCH | 0 | 2 | 7 | 8 | 1 | 0 | 1 | 1 |
| ENT |  | 0 | 1 | 8 | 6 | 2 | 1 | 1 | 0.3411765 |
| FCV |  | 1 | 2 | 9 | 12 | 0 | 0 | 0.2546584 | 1 |
| GND |  | 7 | 1 | 0 | 8 | 0 | 0 | 1 | 1 |
| KAK |  | 1 | 2 | 11 | 11 | 3 | 0 | 0.2177778 | 1 |
| KBO |  | 14 | 1 | 1 | 16 | 0 | 0 | 0.0967742 | 1 |
| KED |  | 15 | 4 | 0 | 15 | 4 | 0 | 1 | 1 |
| KIN' | KIN+BOA | 11 | 10 | 0 | 14 | 6 | 1 | 0.2919408 | 1 |
| KUM |  | 5 | 6 | 8 | 7 | 11 | 1 | 0.1607307 | 0.3449035 |
| KWA |  | 15 | 4 | 0 | 19 | 0 | 0 | 1 | 1 |
| LBV |  | 3 | 5 | 3 | 11 | 0 | 0 | 1 | 1 |
| LPV |  | 11 | 3 | 0 | 14 | 0 | 0 | 1 | 1 |
| MIN |  | 15 | 4 | 0 | 5 | 12 | 2 | 1 | 0.3556129 |
| NGO |  | 7 | 6 | 2 | 13 | 2 | 0 | 1 | 1 |
| OGD' | OGD+GOU+BGR | 1 | 3 | 13 | 14 | 2 | 1 | 0.2883676 | 0.1788856 |
| OHI |  | 9 | 3 | 1 | 6 | 7 | 0 | 0.373913 | 0.4991304 |
| PKT |  | 17 | 1 | 0 | 10 | 7 | 1 | 1 | 1 |
| RAB' | RAB+RABaaf+RABd | 24 | 4 | 0 | 28 | 0 | 0 | 1 | 1 |
| SAN |  | 14 | 0 | 0 | 14 | 0 | 0 | 1 | 1 |
| SHM |  | 7 | 0 | 0 | 7 | 0 | 0 | 1 | 1 |
| THI |  | 4 | 2 | 5 | 6 | 4 | 1 | 0.0610009 | 1 |
| VMB |  | 0 | 2 | 8 | 5 | 5 | 0 | 1 | 1 |

**Table S5.** Association of inversions with chemoreceptor genes (not combined in clusters) and with QTL associated with vector competence in the *Ae. aegypti* genome. Significant associations (P<0.05) are shown in green.

| inv | Or |  |  | Ir |  |  | Gr |  |  |
| --- | --- | --- | --- | --- | --- | --- | --- | --- | --- |
|  | genes count | P | P<sub>adj</sub>(BH) | genes count | P | P<sub>adj</sub>(BH) | genes count | P | P<sub>adj</sub>(BH) |
| 1pA | 0 | 1 | 1 | 0 | 1 | 1 | 0 | 1 | 1 |
| 1pB | 0 | 1 | 1 | 1 | 0.971428377 | 1 | 0 | 1 | 1 |
| 1qC | 0 | 1 | 1 | 0 | 1 | 1 | 0 | 1 | 1 |
| 1qD | 0 | 1 | 1 | 2 | 0.122367965 | 0.508297701 | 0 | 1 | 1 |
| 1qE (Or) | 10 | 0.001919454 | 0.0115167 | 5 | 0.412214825 | 1 | 0 | 1 | 1 |
| 1qF (Or) | 9 | 2.54925·10<sup>-5</sup> | 0.0001967 | 0 | 1 | 1 | 0 | 1 | 1 |
| 1qG | 1 | 0.73875873 | 1 | 1 | 0.785869957 | 1 | 0 | 1 | 1 |
| 2pA (Or) | 10 | 0.001292567 | 0.0087248 | 2 | 0.749540985 | 1 | 0 | 1 | 1 |
| 2qB | 4 | 0.26065767 | 0.7408165 | 4 | 0.140073359 | 0.540282956 | 3 | 0.2096489 | 0.7376148 |
| 2qC | 2 | 0.25152905 | 0.7408165 | 0 | 1 | 1 | 0 | 1 | 1 |
| 2qD | 3 | 0.0270561 | 0.1328209 | 0 | 1 | 1 | 0 | 1 | 1 |
| 3pA | 3 | 0.49502657 | 1 | 16 | 8.96687·10<sup>-7</sup> | 9.68422·10<sup>-6</sup> | 6 | 0.0554054 | 0.2493242 |
| 3pB (Gr) | 7 | 0.34997895 | 0.9449432 | 34 | 2.61432·10<sup>-14</sup> | 4.70578·10<sup>-13</sup> | 31 | 1.937·10<sup>-17</sup> | 1.046·10<sup>-15</sup> |
| 3pC (Ir) | 3 | 0.25382334 | 0.7408165 | 22 | 2.58463·10<sup>-15</sup> | 6.9785·10<sup>-14</sup> | 6 | 0.0094009 | 0.0507646 |
| 3pD (Ir, Gr) | 0 | 1 | 1 | 12 | 2.45061·10<sup>-9</sup> | 3.30832·10<sup>-8</sup> | 7 | 2.177·10<sup>-5</sup> | 0.000196 |
| 3pE | 5 | 0.21855252 | 0.7376148 | 3 | 0.860784011 | 1 | 0 | 1 | 1 |
| 3qF | 0 | 1 | 1 | 0 | 1 | 1 | 0 | 1 | 1 |
| 3qG | 0 | 1 | 1 | 0 | 1 | 1 | 0 | 1 | 1 |

| inv | QTL count | P | P<sub>adj</sub>(BH) |
| --- | --- | --- | --- |
| 1 pA | 1 | 0.18304873 | 0.1830487 |
| 2 pA | 3 | 0.013049448 | 0.0357354 |
| 3 pA | 1 | 0.11875356 | 0.1583381 |
| 3 pB | 2 | 0.017867681 | 0.0357354 |

**Table S6. Chemoreceptor genes combined in clusters.**

Data not shown

**Table S7.** Association of inversions with chemoreceptor gene clusters/genes and with QTL associated with vector competence in the *Ae. aegypti* genome. Significant associations (P<0.1) are shown in green.

| inv | Or |  |  | Ir |  |  | Gr |  |  |
| --- | --- | --- | --- | --- | --- | --- | --- | --- | --- |
|  | genes count | P | P<sub>adj</sub>(BH) | genes count | P | P<sub>adj</sub>(BH) | genes count | P | P<sub>adj</sub>(BH) |
| 1pA | 0 | 1 | 1 | 0 | 1 | 1 | 0 | 1 | 1 |
| 1pB | 0 | 1 | 1 | 1 | 0.6823751 | 1 | 0 | 1 | 1 |
| 1qC | 0 | 1 | 1 | 0 | 1 | 1 | 0 | 1 | 1 |
| 1qD | 0 | 1 | 1 | 2 | 0.0154628 | 0.1391653 | 0 | 1 | 1 |
| 1qE | 6 | 0.0146183 | 0.13916529 | 3 | 0.143392 | 0.5956282 | 0 | 1 | 1 |
| 1qF (Or) | 5 | 0.0023926 | 0.064600713 | 0 | 1 | 1 | 0 | 1 | 1 |
| 1qG | 1 | 0.548619 | 1 | 1 | 0.3917396 | 1 | 0 | 1 | 1 |
| 2pA | 5 | 0.057149 | 0.34289384 | 2 | 0.5102658 | 1 | 0 | 1 | 1 |
| 2qB | 2 | 0.5024941 | 1 | 4 | 0.0367536 | 0.2754 | 3 | 0.1041584 | 0.4687128 |
| 2qC | 1 | 0.4650474 | 1 | 0 | 1 | 1 | 0 | 1 | 1 |
| 2qD | 3 | 0.0079785 | 0.1077104 | 0 | 1 | 1 | 0 | 1 | 1 |
| 3pA | 2 | 0.5740006 | 1 | 3 | 0.0820928 | 0.403001 | 1 | 0.6585495 | 1 |
| 3pB (Gr) | 6 | 0.2271446 | 0.87612933 | 5 | 0.0660943 | 0.3569094 | 9 | 1.19E-04 | 6.43·10<sup>-3</sup> |
| 3pC (Ir) | 2 | 0.3606952 | 1 | 4 | 0.0042375 | 0.0762744 | 1 | 0.5059732 | 1 |
| 3pD | 0 | 1 | 1 | 1 | 0.2787067 | 1 | 2 | 4.08E-02 | 0.2754 |
| 3pE | 2 | 0.6868696 | 1 | 1 | 0.7344877 | 1 | 0 | 1 | 1 |
| 3qF | 0 | 1 | 1 | 0 | 1 | 1 | 0 | 1 | 1 |
| 3qG | 0 | 1 | 1 | 0 | 1 | 1 | 0 | 1 | 1 |

**Table S8.** Breakpoint coordinates for *An. gambiae* inversions. Telomeric and centromeric breakpoint information are provided for 2Rj (Coulibaly *et al*. 2007), 2Rb (Lobo *et al*. 2010), 2Rc, 2Rd, 2Ru, 2Rb (George *et al*. 2010), and 2La (Sharakhov *et al*. 2006).

| Name | Telomeric breakpoint | Centromeric breakpoint | Reference |
| --- | --- | --- | --- |
| 2Rj | 3,262,186 bp | 15,750,717 bp | Colibaly *et al.*, 2007 |
| 2Rb | 19023925 bp | 26758676 bp | Lobo *et al.*, 2010 |
| 2Rc | 26.78 Mb | 31.45 Mb | George *et al.*, 2010 |
| 2Rd | 31.48 Mb | 42.6 Mb | George *et al.*, 2010 |
| 2Ru | 31.48 Mb | 35.9 Mb | George *et al.*, 2010 |
| 2Rbk | 25.1 Mb | 37.55 Mb | George *et al.*, 2010 |
| 2La | 20,524,058 bp | 42,165,532 bp | Sharakhov *et al.*, 2006 |

## References

WHO, 2022. Dengue and severe dengue. https://www.who.int/news-room/fact-sheets/detail/den-gue-and-severe-dengue. Accessed 22 Jan 2024.

Andrews, S., 2010 FastQC: a quality control tool for high throughput sequnce data. https://www.bioinformat-ics.babraham.ac.uk/projects/fastqc/. Accessed 22 Jan 2024.

Arensburger, P., K. Megy, R. M. Waterhouse, J. Abrudan, P. Amedeo et al., 2010 Sequencing of Culex quinquefasciatus establishes a platform for mosquito comparative genomics. Science 330: 86–88.

Artemov, G. N., V. S. Fedorova, D. A. Karagodin, Brusentsov, II, E. M. Baricheva et al., 2021 New cytogenetic photomap and molecular diagnostics for the cryptic species of the malaria mosquitoes Anopheles messeae and Anopheles daciae from Eurasia. Insects 12:1–16.

Aubry, F., S. Dabo, C. Manet, I. Filipovic, N. H. Rose et al., 2020 Enhanced Zika virus susceptibility of globally invasive Aedes aegypti populations. Science 370: 991–996.

Ayala, D., P. Acevedo, M. Pombi, I. Dia, D. Boccolini et al., 2017 Chromosome inversions and ecological plasticity in the main African malaria mosquitoes. Evolution 71: 686–701.

Ayala, D., A. Ullastres and J. Gonzalez, 2014 Adaptation through chromosomal inversions in Anopheles. Front Genet 5: 129.

Ayala, F. J., and M. Coluzzi, 2005 Chromosome speciation: humans, Drosophila, and mosquitoes. Proc Natl Acad Sci U S A 102 Suppl 1: 6535–6542.

Benedict, M., and E. M. Dotson, 2015 Methods in Anopheles research. pp408.

Benjamini, Y., and Y. Hochberg, 1995 Controlling the false discovery rate: a practical and powerful approach to multiple testing. Journal of the Royal Statistical Society: Series B (Methodological) 57: 129–300.

Berdan, E. L., N. H. Barton, R. Butlin, B. Charlesworth, R. Faria et al., 2023 How chromosomal inversions reorient the evolutionary process. J Evol Biol. 36(12): 1761–1782.

Berdan, E. L., T. Flatt, G. M. Kozak, K. E. Lotterhos and B. Wielstra, 2022 Genomic architecture of supergenes: connecting form and function. Philos Trans R Soc Lond B Biol Sci 377: 20210192.

Bernhardt, S. A., C. Blair, M. Sylla, C. Bosio and W. C. t. Black, 2009 Evidence of multiple chromosomal inversions in Aedes aegypti formosus from Senegal. Insect Mol Biol 18: 557–569.

Bosio, C. F., R. E. Fulton, M. L. Salasek, B. J. Beaty and W. C. t. Black, 2000 Quantitative trait loci that control vector competence for dengue-2 virus in the mosquito Aedes aegypti. Genetics 156: 687–698.

Brown, J. E., B. R. Evans, W. Zheng, V. Obas, L. Barrera-Martinez et al., 2014 Human impacts have shaped historical and recent evolution in Aedes aegypti, the dengue and yellow fever mosquito. Evolution 68: 514–525.

Brown, J. E., C. S. McBride, P. Johnson, S. Ritchie, C. Paupy et al., 2011 Worldwide patterns of genetic differentiation imply multiple ‘domestications’ of Aedes aegypti, a major vector of human diseases. Proc Biol Sci 278: 2446–2454.

Brusentsov, II, M. I. Gordeev, A. A. Yurchenko, D. A. Karagodin, A. V. Moskaev et al., 2023 Patterns of genetic differentiation imply distinct phylogeographic history of the mosquito species Anopheles messeae and Anopheles daciae in Eurasia. Mol Ecol 32: 5609–5625.

Cabrera, C. P., P. Navarro, J. E. Huffman, A. F. Wright, C. Hayward et al., 2012 Uncovering networks from genome-wide association studies via circular genomic permutation. G3 (Bethesda) 2: 1067–1075.

Campos, J., C. F. Andrade and S. M. Recco-Pimentel, 2003 A technique for preparing polytene chromosomes from Aedes aegypti (Diptera, Culicinae). Mem Inst Oswaldo Cruz 98: 387–390.

Chakraborty, A., and F. Ay, 2017 Identification of copy number variations and translocations in cancer cells from Hi-C data. Bioinformatics. 34(2):338–345.

Coluzzi, M., A. Sabatini, A. della Torre, M. A. Di Deco and V. Petrarca, 2002 A polytene chromosome analysis of the Anopheles gambiae species complex. Science 298: 1415–1418.

Corbett-Detig, R. B., I. Said, M. Calzetta, M. Genetti, J. McBroome et al., 2019 Fine-mapping complex inversion breakpoints and investigating somatic pairing in the Anopheles gambiae species complex using proximity-ligation sequencing. Genetics 213: 1495–1511.

Coulibaly, M. B., N. F. Lobo, M. C. Fitzpatrick, M. Kern, O. Grushko et al., 2007 Segmental duplication implicated in the genesis of inversion 2Rj of Anopheles gambiae. PLoS One 2: e849.

Dekker, J., K. Rippe, M. Dekker and N. Kleckner, 2002 Capturing chromosome conformation. Science 295: 1306–1311.

Dickson, L. B., I. Sanchez-Vargas, M. Sylla, K. Fleming and W. C. . Black, 2014 Vector competence in West African Aedes aegypti Is Flavivirus species and genotype dependent. PLoS Negl Trop Dis 8: e3153.

Dickson, L. B., M. V. Sharakhova, V. A. Timoshevskiy, K. L. Fleming, A. Caspary et al., 2016 Reproductive incompatibility involving Senegalese Aedes aegypti (L) is associated with chromosome rearrangements. PLoS Negl Trop Dis 10: e0004626.

Durand, N. C., J. T. Robinson, M. S. Shamim, I. Machol, J. P. Mesirov et al., 2016 Juicebox provides a visualization system for Hi-C contact maps with unlimited zoom. Cell Syst 3: 99–101.

Emms, D. M., and S. Kelly, 2019 OrthoFinder: phylogenetic orthology inference for comparative genomics. Genome Biol 20: 238.

Engreitz, J. M., V. Agarwala and L. A. Mirny, 2012 Three-dimensional genome architecture influences partner selection for chromosomal translocations in human disease. PLoS One 7: e44196.

Faria, N. R., S. Azevedo Rdo, M. U. Kraemer, R. Souza, M. S. Cunha et al., 2016 Zika virus in the Americas: Early epidemiological and genetic findings. Science 352: 345–349.

Ferguson-Smith, M. A., and V. Trifonov, 2007 Mammalian karyotype evolution. Nat Rev Genet 8: 950–962.

Fijman, N. S., and D. A. Yee, 2022 Mapping Yellow fever epidemics as a potential indicator of the historical range of Aedes aegypti in the United States. Mem Inst Oswaldo Cruz 117: e220306.

Fontenille, D., and J. R. Powell, 2020 From anonymous to public enemy: how does a mosquito become a feared arbovirus vector? Pathogens 9(4):1–11.

Fuller, Z. L., S. A. Koury, N. Phadnis and S. W. Schaeffer, 2019 How chromosomal rearrangements shape adaptation and speciation: Case studies in Drosophila pseudoobscura and its sibling species Drosophila persimilis. Mol Ecol 28: 1283–1301.

Garcia-Luna, S. M., J. Weger-Lucarelli, C. Ruckert, R. A. Murrieta, M. C. Young et al., 2018 Variation in competence for ZIKV transmission by Aedes aegypti and Aedes albopictus in Mexico. PLoS Negl Trop Dis 12: e0006599.

George, P., M. V. Sharakhova and I. V. Sharakhov, 2010 High-resolution cytogenetic map for the African malaria vector Anopheles gambiae. Insect Mol Biol 19: 675–682.

Gloria-Soria, A., D. Ayala, A. Bheecarry, O. Calderon-Arguedas, D. D. Chadee et al., 2016 Global genetic diversity of Aedes aegypti. Mol Ecol 25: 5377–5395.

Gray, E. M., K. A. Rocca, C. Costantini and N. J. Besansky, 2009 Inversion 2La is associated with enhanced desiccation resistance in Anopheles gambiae. Malar J 8: 215.

Gubler, D. J., 2012 The 20th century re-emergence of epidemic infectious diseases: lessons learned and future prospects. Med J Aust 196: 293–294.

Hall, A. B., S. Basu, X. Jiang, Y. Qi, V. A. Timoshevskiy et al., 2015 SEX DETERMINATION. A male-determining factor in the mosquito Aedes aegypti. Science 348: 1268–1270.

Harewood, L., K. Kishore, M. D. Eldridge, S. Wingett, D. Pearson et al., 2017 Hi-C as a tool for precise detection and characterisation of chromosomal rearrangements and copy number variation in human tumours. Genome Biol 18: 125.

Holt, R. A., G. M. Subramanian, A. Halpern, G. G. Sutton, R. Charlab et al., 2002 The genome sequence of the malaria mosquito Anopheles gambiae. Science 298: 129–149.

Huang, K., and L. H. Rieseberg, 2020 Frequency, origins, and evolutionary role of chromosomal inversions in plants. Front Plant Sci 11: 296.

Jay, P., and M. Joron, 2022 The double game of chromosomal inversions in a neotropical butterfly. C R Biol 345: 57–73.

Kitzmiller, J. B., 1976 Genetics, cytogenetics, and evolution of mosquitoes., pp. 315–433 in Advances in Genetics, edited by E. W. Caspari. Elsevier Inc.

Kosuthova, K., and R. Solc, 2023 Inversions on human chromosomes. Am J Med Genet A 191: 672–683.

Kotsakiozi, P., B. R. Evans, A. Gloria-Soria, B. Kamgang, M. Mayanja et al., 2018 Population structure of a vector of human diseases: Aedes aegypti in its ancestral range, Africa. Ecol Evol 8: 7835–7848.

Krimbas, C. B., and J. R. Powell, 1992 Drosophila inversion polymorphism. CRC Press. 560 pp.

Krzywinski, M., J. Schein, I. Birol, J. Connors, R. Gascoyne et al., 2009 Circos: an information aesthetic for comparative genomics. Genome Res 19: 1639–1645.

Lanzaro, G. C., Y. T. Toure, J. Carnahan, L. Zheng, G. Dolo et al., 1998 Complexities in the genetic structure of Anopheles gambiae populations in west Africa as revealed by microsatellite DNA analysis. Proc Natl Acad Sci U S A 95: 14260–14265.

Liang, J., S. M. Bondarenko, I. V. Sharakhov and M. V. Sharakhova, 2022 Obtaining polytene, meiotic, and mitotic chromosomes from mosquitoes for cytogenetic analysis. Cold Spring Harb Protoc.

Lieberman-Aiden, E., N. L. van Berkum, L. Williams, M. Imakaev, T. Ragoczy et al., 2009 Comprehensive mapping of long-range interactions reveals folding principles of the human genome. Science 326: 289–293.

Lobo, N. F., D. M. Sangare, A. A. Regier, K. R. Reidenbach, D. A. B etz et al., 2010 Breakpoint structure of the Anopheles gambiae 2Rb chromosomal inversion. Malar J 9: 293.

Lorenz, L., B. J. Beaty, T. H. Aitken, G. P. Wallis and W. J. Tabachnick, 1984 The effect of colonization upon Aedes aegypti susceptibility to oral infection with yellow fever virus. Am J Trop Med Hyg 33: 690–694.

Love, R. R., M. Pombi, M. W. Guelbeogo, N. R. Campbell, M. T. Stephens et al., 2020 Inversion genotyping in the Anopheles gambiae complex using high-throughput array and sequencing platforms. G3 (Bethesda) 10: 3299–3307.

Love, R. R., S. N. Redmond, M. Pombi, B. Caputo, V. Petrarca et al., 2019 In silico karyotyping of chromosomally polymorphic malaria mosquitoes in the Anopheles gambiae complex. G3 (Bethesda) 9: 3249–3262.

Love, R. R., A. M. Steele, M. B. Coulibaly, S. F. Traore, S. J. Em ich et al., 2016 Chromosomal inversions and ecotypic differentiation in Anopheles gambiae: the perspective from whole-genome sequencing. Mol Ecol 25: 5889–5906.

Lukyanchikova, V., V. Fishman and I. Sharakhov, 2022a In situ Hi-C for mosquito embryos. Protocol Exchange. pp. 1–11.

Lukyanchikova, V., M. Nuriddinov, P. Belokopytova, A. Taskina, J. Liang et al., 2022b Anopheles mosquitoes reveal new principles of 3D genome organization in insects. Nat Commun 13: 1960.

Masri, R. A., D. A. Karagodin, A. Sharma and M. V. Sharakhova, 2021 A gene-based method for cytogenetic mapping of repeat-rich mosquito genomes. Insects 12:1–11.

Matthews, B. J., O. Dudchenko, S. B. Kingan, S. Koren, I. Antoshechkin et al., 2018 Improved reference genome of Aedes aegypti informs arbovirus vector control. Nature 563: 501–507.

Mattingly, P. F., 1957 Genetical aspects of the Aedes aegypti problem. I. Taxonomy and bionomics. Ann Trop Med Parasitol 51: 392–408.

McBride, C. S., 2016 Genes and odors underlying the recent evolution of mosquito preference for humans. Curr Biol 26: R41–46.

McBride, C. S., F. Baier, A. B. Omondi, S. A. Spitzer, J. Lutomiah et al., 2014 Evolution of mosquito preference for humans linked to an odorant receptor. Nature 515: 222–227.

Metz, H. C., A. K. Miller, J. You, J. Akorli, F. W. Avila et al., 2023 Evolution of a mosquito’s hatching behavior to match its human-provided habitat. Am Nat 201: 200–214.

Nabel, G. J., and E. A. Zerhouni, 2016 Once and future epidemics: Zika virus emerging. Sci Transl Med 8: 330ed332.

Olagnier, D., D. Amatore, L. Castiello, M. Ferrari, E. Palermo et al., 2016 Dengue virus immunopathogenesis: lessons applicable to the emergence of Zika virus. J Mol Biol 428(17): 3429–3448.

Oliveira da Silva, W., S. M. Malcher, M. A. Ferguson-Smith, P. C. M. O’Brien, R. V. Rossi et al., 2024 Chromosomal rearrangements played an important role in the speciation of rice rats of genus Cerradomys (Rodentia, Sigmodontinae, Oryzomyini). Sci Rep 14: 545.

Powell, J. R., 2016a Mosquitoes on the move. Science 354: 971–972.

Powell, J. R., 2016b Mosquitoes: New contender for most lethal animal. Nature 540: 525.

Powell, J. R., A. Gloria-Soria and P. Kotsakiozi, 2018 Recent history of Aedes aegypti: vector genomics and epidemiology records. Bioscience 68: 854–860.

Powell, J. R., and W. J. Tabachnick, 2013 History of domestication and spread of Aedes aegypti--a review. Mem Inst Oswaldo Cruz 108 Suppl 1: 11–17.

Quinlan, A. R., 2014 BEDTools: the Swiss-army tool for genome feature analysis. Curr Protoc Bioinformatics 47:11.12.1–11.12.34.

R_Core_Team, 2021 R: A language and environment for statistical computing. Vienna: R foundation for statistical computing. https://www.r-project.org/. Accessed 22 Jan 2024.

Redmond, S. N., A. Sharma, I. Sharakhov, Z. Tu, M. Sharakhova et al., 2020 Linked-read sequencing identifies abundant microinversions and introgression in the arboviral vector Aedes aegypti. BMC Biol 18: 26.

Riehle, M. M., T. Bukhari, A. Gneme, W. M. Guelbeogo, B. Coulibaly et al., 2017 The Anopheles gambiae 2La chromosome inversion is associated with susceptibility to Plasmodium falciparum in Africa. Elife 6:1–24.

Robinson, J. T., D. Turner, N. C. Durand, H. Thorvaldsdottir, J. P. Mesirov et al., 2018 Juicebox.js provides a cloud-based visualization system for Hi-C data. Cell Syst 6: 256–258 e251.

Rose, N. H., A. Badolo, M. Sylla, J. Akorli, S. Otoo et al., 2023 Dating the origin and spread of specialization on human hosts in Aedes aegypti mosquitoes. Elife 12:1–17.

Rose, N. H., M. Sylla, A. Badolo, J. Lutomiah, D. Ayala et al., 2020 Climate and urbanization drive mosquito preference for humans. Curr Biol 30: 3570–3579 e3576.

Ryazansky, S. S. et al., 2024 The chromosome-scale genome assembly for the West Nile vector Culex quinquefasciatus uncovers patterns of genome evolution in mosquitoes. BMC Biology 22:16.

Severson, D. W., J. K. Meece, D. D. Lovin, G. Saha and I. Morlais, 2002 Linkage map organization of expressed sequence tags and sequence tagged sites in the mosquito, Aedes aegypti. Insect Mol Biol 11: 371–378.

Sharakhov, I. V., B. J. White, M. V. Sharakhova, J. Kayondo, N. F. Lobo et al., 2006 Breakpoint structure reveals the unique origin of an interspecific chromosomal inversion (2La) in the Anopheles gambiae complex. Proc Natl Acad Sci U S A 103: 6258–6262.

Sharakhova, M. V., G. N. Artemov, V. A. Timoshevskiy and I. V. Sharakhov, 2019 Physical genome mapping using fluorescence in situ hybridization with mosquito chromosomes. Methods Mol Biol 1858: 177–194.

Sharakhova, M. V., V. A. Timoshevskiy, F. Yang, S. Demin, D. W. Severson et al., 2011a Imaginal discs: a new source of chromosomes for genome mapping of the yellow fever mosquito Aedes aegypti. PLoS Negl Trop Dis 5: e1335.

Sharakhova, M. V., A. Xia, S. C. Leman and I. V. Sharakhov, 2011b Arm-specific dynamics of chromosome evolution in malaria mosquitoes. BMC Evol Biol 11: 91.

Shenlab-sinai, 2023 GeneOverlap: Test and visualize gene overlaps. R package version 1.36.0. http://shen-lab-sinai.github.io/shenlab-sinai/. Accessed 22 Jan 2024.

Shimodaira, H., 2002 An approximately unbiased test of phylogenetic tree selection. Syst Biol 51: 492–508.

Shimodaira, H., 2004 Approximately unbiased tests of regions using multistep-multiscale bootstrap resampling. Annals of Statistics: 2616–2641.

Simard, F., D. Ayala, G. C. Kamdem, M. Pombi, J. Etouna et al., 2009 Ecological niche partitioning between Anopheles gambiae molecular forms in Cameroon: the ecological side of speciation. BMC Ecol 9: 17.

Soghigian, J., A. Gloria-Soria, V. Robert, G. Le Goff, A. B. Failloux et al., 2020 Genetic evidence for the origin of Aedes aegypti, the yellow fever mosquito, in the southwestern Indian Ocean. Mol Ecol 29: 3593–3606.

Souza-Neto, J. A., J. R. Powell and M. Bonizzoni, 2019 Aedes aegypti vector competence studies: A review. Infect Genet Evol 67: 191–209.

Stegniy, V. N., 1991 Population genetics and evolution of malaria mosquitoes. Tomsk State University Publisher.

Suzuki, R., and H. Shimodaira, 2006 Pvclust: an R package for assessing the uncertainty in hierarchical clustering. Bioinformatics 22: 1540–1542.

Sylla, M., C. Bosio, L. Urdaneta-Marquez, M. Ndiaye and W. C. t. Black, 2009 Gene flow, subspecies composition, and dengue virus-2 susceptibility among Aedes aegypti collections in Senegal. PLoS Negl Trop Dis 3: e408.

Tabachnick, W. J., 2013 Nature, nurture and evolution of intra-species variation in mosquito arbovirus transmission competence. Int J Environ Res Public Health 10: 249–277.

Tabachnick, W. J., G. P. Wallis, T. H. Aitken, B. R. Miller, G. D. Amato et al., 1985 Oral infection of Aedes aegypti with yellow fever virus: geographic variation and genetic considerations. Am J Trop Med Hyg 34: 1219–1224.

Thompson, M. J., and C. D. Jiggins, 2014 Supergenes and their role in evolution. Heredity (Edinb) 113: 1–8.

Timoshevskiy, V. A., N. A. Kinney, B. S. deBruyn, C. Mao, Z. Tu et al., 2014 Genomic composition and evolution of Aedes aegypti chromosomes revealed by the analysis of physically mapped supercontigs. BMC Biol 12: 27.

Timoshevskiy, V. A., D. W. Severson, B. S. Debruyn, W. C. Black, I. V. Sharakhov et al., 2013 An integrated linkage, chromosome, and genome map for the yellow fever mosquito Aedes aegypti. PLoS Negl Trop Dis 7: e2052.

Timoshevskiy, V. A., A. Sharma, I. V. Sharakhov and M. V. Sharakhova, 2012 Fluorescent in situ hybridization on mitotic chromosomes of mosquitoes. J Vis Exp. 67:1–7.

van Berkum, N. L., E. Lieberman-Aiden, L. Williams, M. Imakaev, A. Gnirke et al., 2010 Hi-C: a method to study the three-dimensional architecture of genomes. J Vis Exp. 39:1–7.

Wallis, G. P., T. H. Aitken, B. J. Beaty, L. Lorenz, G. D. Amato et al., 1985 Selection for susceptibility and refractoriness of Aedes aegypti to oral infection with yellow fever virus. Am J Trop Med Hyg 34: 1225–1231.

Weaver, S. C., F. Costa, M. A. Garcia-Blanco, A. I. Ko, G. S. Ribeiro et al., 2016 Zika virus: history, emergence, biology, and prospects for control. Antiviral Res 130: 69–80.

Westram, A. M., R. Faria, K. Johannesson, R. Butlin and N. Barton, 2022 Inversions and parallel evolution. Philos Trans R Soc Lond B Biol Sci 377: 20210203.

Wolfram, I., 2023 Mathematica. The world’s definitive system for modern technical computing. Champaign, Illinois, Wolfram Research Inc. https://www.wolfram.com/mathematica, Accessed 22 Jan 2024.

